# Newly Acquired Word-Action Associations Trigger Auditory Cortex Activation During Movement Preparation: Implications for Hebbian Plasticity in Action Words Learning

**DOI:** 10.1101/2024.09.05.611409

**Authors:** V.D. Tretyakova, A.A. Pavlova, V.V. Arapov, A.M. Rytikova, A.U. Nikolayeva, A.O. Prokofyev, B.V. Chernyshev, T.A. Stroganova

## Abstract

Hebbian learning is believed to play a key role in acquisition of action words. However, this biological mechanism requires activation of the neural assemblies representing a word form and a corresponding movement to repeatedly overlap in time. In reality, though, these associated events could be separated by seconds. In the current MEG study, we examined trial-and-error learning of associations between novel auditory pseudowords and movements of specific body parts. We aimed to explore how the brain bridges the temporal gap between the transient activity evoked by auditory input and the preparatory motor activation before the corresponding movement. To address this, we compared learning-induced changes in neuromagnetic responses locked to the onset of the stimulus and to the onset of the movement. As learning progressed, both types of neural responses showed sustained enhancement during the delay period between the auditory pseudoword and the required movement. Cortical sources of this learning-induced increase were localized bilaterally in the lateral and medial temporal cortices. Notably, the learning effect was significantly stronger when measured time-locked to the movement onset, rather than to the pseudoword onset. This suggests that, after the pseudoword-movement associations were reliably acquired, the non-primary auditory cortex was reactivated in sync with the preparation of the upcoming movement. Such reactivation likely served to bring together in time the representations of the correct action and the preceding auditory cue. This temporal alignment could enable Hebbian learning, leading to long-lasting synaptic changes in temporally correlated neural assemblies.

## Introduction

It is generally accepted that language acquisition involves biological mechanisms of associative learning (Barsalou, 2003; Pulvermüller, 2005, 2018). These theories crucially rely on the principle of activity-dependent modification of synaptic strengths triggered by repeated co-activation of two neuronal representations (Hebb, 1949; Nicoll, 2017). For example, according to this view, meaning of action words is acquired because neural assemblies coding the auditory representation of the word and neurons programming the corresponding physical movement are repeatedly activated simultaneously, which eventually leads to strengthened synaptic connections and formation of distributed and interrelated ‘cell assemblies’ (Hebb, 1949). As a result of such plastic changes, the auditory signal presented alone begins to activate both neuronal populations – not only the auditory areas, but the motor regions as well.

Indirect evidence for applicability of Hebbian learning principle to human language was mainly derived from the neuroimaging findings on simultaneous activation in both auditory and somatotopically organized motor cortical areas driven by auditory presentations of well-learnt action words (Shtyrov, Butorina, Nikolaeva, & Stroganova, 2014). Furthermore, this principle was supported by computational modeling of action word learning (Tomasello, Garagnani, Wennekers, & Pulvermüller, 2017).

However, biologically-based concepts of action word learning have important assumption that, throughout the learning procedure, the auditory activation produced by an action word and programming of corresponding motor commands in the motor cortex should overlap in time or follow each other with a very small delay (see e.g., Pulvermüller, 2018). Particularly, studies of spike timing dependent plasticity (STDP) – a biological mechanism which is often identified with Hebbian learning – have defined that time windows between pre- and post-synaptic inputs could not exceed few tens of milliseconds to STDP takes place (Bi & Poo, 1998; Dan & Poo, 2006; Egger, Feldmeyer, & Sakmann, 1999; Koch, Ponzo, Di Lorenzo, Caltagirone, & Veniero, 2013). However, in a natural environment such learning can occur despite a temporal gap between an action and an associated word on a timescale of hundreds or even thousands of milliseconds. To explain action word learning by spike-timing-dependent plasticity, as it was proposed by Pulvermuller (2018), one must account for this large discrepancy in timescales, and find a mechanism that bridges up this gap in time.

Available neuronal studies on animals have suggested two possible mechanisms that could allow learning to occur in the adult brain, despite the lack of precise temporal coincidence between the sensory signal and the motor response. One approach to this problem is to assume that along the course of associative learning, novel action words start to generate persistent auditory responses that outlast these stimuli long enough to overlap with the associated motor command. Since Hebb’s idea (Hebb, 1949) that was put forward more than 50 years ago, mnemonic activity has been hypothesized to be sustained by synaptic reverberation in a recurrent circuit (Drew & Abbott, 2006; Shu, Hasenstaub, & McCormick, 2003). We will address such presumable prolongation of activity triggered by stimulus onset as a “retrospective” mechanism. This possibility was supported by numerous findings of increased neural activity during the delay period in working memory tasks in monkeys (X.-J. Wang, 2001) and humans (Vogel & Machizawa, 2004).

Another option is that the input representation could be “prospectively” re-activated shortly before the movement onset so that the two populations of auditory and motor neurons would be activated concurrently. This view is in line with proposals that stimulus-related activity can subside during the delay and then reappear when a task requires (Lewis-Peacock, Drysdale, Oberauer, & Postle, 2012; Masse, Rosen, & Freedman, 2020). Similarly, in an associative learning task where participants were acquiring associations between short videos and unrelated nouns, “reactivation” of the video-specific neural response in EEG signal was shown during delayed presentation of the noun associate (Michelmann, Bowman, & Hanslmayr, 2018). However, usage of non-phased locked activity in that study did not allow to conclude whether the video-specific pattern reflected maintenance of the stimulus-related activity over the delay period or it was ignited anew by the noun presentation.

Whereas the two models are not mutually exclusive – and both mechanisms can coexist and provide cumulative effects – the critical difference between them is timing of the auditory activation during the delay period between presentation of a novel action word and a chosen motor response. The “retrospective” model predicts that, over the course of learning, a rapidly decaying auditory neural response will be replaced by the more persistent auditory activation that, although decreasing over time, will still last until the end of the delay period. In contrast, if the co-occurrence of auditory and motor activation during the delay period is driven by “prospective” re-activation of the former representation, its neural manifestations in the form of increased auditory activation will be strongest immediately before the motor act.

Our recent magnetoencephalography (MEG) study in human adults indicated that passive listening to pseudowords previously associated with specific motor actions through active operant conditioning evokes differential brain activation in auditory and other speech areas of the left hemisphere compared with pseudowords without motor associations (Razorenova et al., 2020). The emergent difference in neural responsiveness that persisted throughout the passive listening of an auditory cue proved that active learning of auditory-action association drives rapid cortical plasticity in auditory cortex of human adults, and that finding was a necessary prerequisite for the current study.

Here, we investigated the learning-induced neural changes during active formation of pseudoword-action association in the same subjects. For this study, we applied a set of eight tightly controlled action-related and non-action-related pseudowords, whose relatedness to action of different body parts was gradually established by the participants through trial-and-error learning.

First, we aimed to investigate if the temporal gap between an auditory pseudoword and a motor action is indeed filled with the persistent activation of auditory neural representations. For this purpose, we compared brain activity in response to the auditory pseudowords between early and advanced stages of learning. We expected that, as the participants shifted from random guessing to purposeful responding according to emerging understanding of the associative rules, neural response to the auditory pseudowords would become strengthened and prolonged to ensure co-occurrence and, thus, association of the auditory activation with the preparation of the chosen motor program.

If this persistent activation was present during the delay period, we aimed to probe which of the two suggested neural accounts – “retrospective” or “prospective” – better explains this result. We planned to compare cortical responses time-locked to the auditory stimulus onset and to the motor response onset. If the “retrospective” hypothesis is correct, we could expect a learning-induced prolongation of auditory cue-evoked activation which would be the most evident in the stimulus-locked data. The “prospective” hypothesis predicts that the advanced learning stage would be mainly characterized by the appearance of the auditory activation that, paradoxically, would be largely driven by preparation for motor action. Hence, the auditory activation would be time-locked to the onset of the motor response, and would co-occur with activation of the premotor areas that form a central node of action preparation.

## Materials and Methods

### Participants

Twenty-eight native Russian speakers (mean age 25.5 years, range 19-38 years, 17 males) volunteered to participate in the study. The number of participants aligns with previous M/EEG studies that examined protracted ERF/ERP components during the delay period in working memory tasks (Fan, Muthukumaraswamy, Singh, & Shapiro, 2012; Günseli et al., 2019; Jafarpour, Penny, Barnes, Knight, & Duzel, 2017; Kuo, Nobre, Scerif, & Astle, 2016). All participants were right-handed (Edinburgh Handedness Inventory (Oldfield, 1971)); they had normal hearing and reported no history of neurological or psychiatric disorders. The study was conducted following the ethical principles regarding human experimentation (Helsinki Declaration) and approved by the Ethics Committee of the Moscow State University of Psychology and Education. All participants signed an informed consent before the experiment.

### Stimuli and Procedure

The auditory stimuli (pseudowords) were designed so that their acoustic and phonetic properties were controlled and balanced while their lexical status was manipulated by learning (for their detailed description see Razorenova et al., 2020). We used nine consonant-vowel (CV) syllables, which were organized in eight disyllabic (C_1_V_1_ C_2_V_2_) novel meaningless word-forms (pseudowords). The resulting pseudowords complied with Russian language phonetics and phonotactic constraints. During the associative learning procedure, four of these word stimuli were associated with a unique action performed by one of four body extremities (action-related pseudowords, APW), while the other four were not assigned any motor response (non-action-related pseudowords, NPW).

The first two phonemes (C_1_V_1_) formed the syllable ‘hi’ (xʲˈi) that was identical for all pseudowords used. The next two phonemes (C_2_ and V_2_) were balanced across APW and NPW stimuli in such a way to ensure that acoustic and phonetic features were fully matched between the two stimuli types (Table 1). The third phonemes (C_2_) – one of the four consonants ‘ch’ (^t͡^ɕ), ‘sh’ (ʂ), ‘s’ (^s̪^), ‘v’ (v) – occurred in both APW and NPW stimuli and signaled which extremity a subject might be prepared to use (right hand, left hand, right foot, or left foot). The fourth phonemes (V_2_: vowel ‘a’ (ə) or ‘u’ (ʊ)) were counterbalanced between APW and NPW conditions being included in the two stimuli of each type so that to form eight unique phonemic combinations. Thus, only the fourth phoneme allows recognition of all pseudo-words used in the experiment. Onset of the fourth phoneme henceforth will be referred to as “word-form uniqueness point”.

**Table 1.**
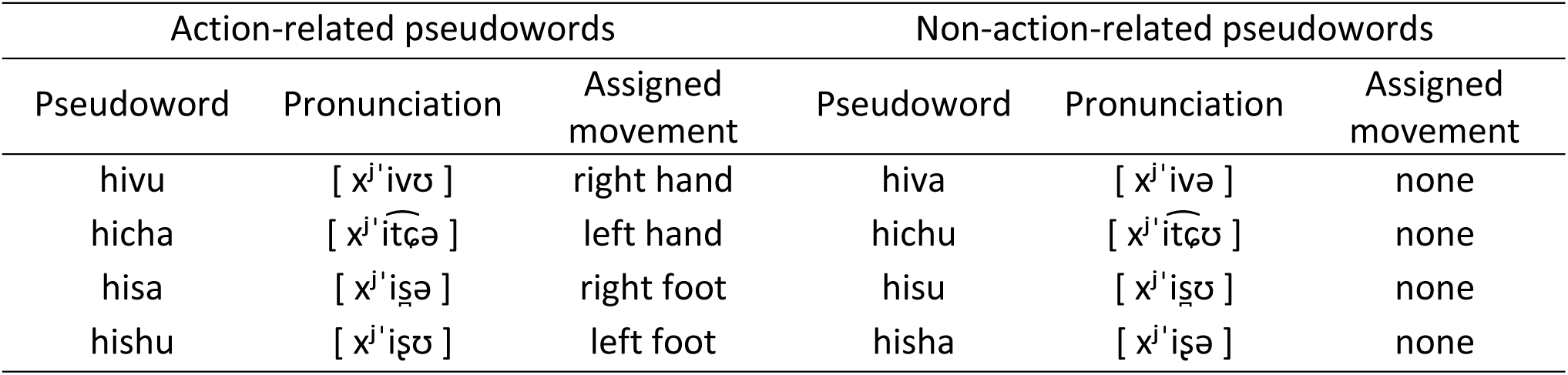
Stimulus-to-response mapping.

As can be seen in Table 1, the phonetic composition of the stimuli and stimulus-to-response mapping complied with a full within-subject counterbalanced design, in relation to the third and fourth phonemes, as well as in relation to movements by left/right and upper/lower extremities.

All stimuli were digitally recorded (PCM, 32 bit, 22050 Hz, 1 channel, 352 kbps) by a female native Russian speaker’s voice in a sound-attenuated booth. We generated eight pseudoword stimuli by way of cross-splicing the sound recordings. We used a single variant of the first syllable (C_1_V_1_), four variants of the third phoneme (C_2_) and two variants of the last vowel (V_2_). All pseudowords were pronounced with the first vowel ‘i’ stressed. The amplitude of the sound recordings was digitally equalized by maximal power, which corresponded to the stressed vowel ‘i’. Cross-splicing and normalization of the recorded stimuli was done using Adobe Audition CS6.5 software. The mean duration of the pseudowords was approximately 480 ms, with word-form uniqueness point occurring approximately 360 ms after the stimulus onset.

Two additional non-speech sounds were used as positive and negative feedback signals, each 400 ms in length. The positive and negative stimuli differed in their spectral frequency (ranges were approximately 400–800 Hz for positive and 65–100 Hz for negative feedback), with spectral maxima increasing in frequency over time for the positive feedback and decreasing for the negative feedback.

Hand responses (Table 1) were recorded using hand-held buttons (package 932, CurrentDesigns, Philadelphia, PA, USA) pressed by the right or left thumb while foot movements were recorded by means of custom-made pedals pushed by the right or left foot. In order to minimize movement artifacts, we kept the trajectory of the movements rather short (<1 cm for buttons and <3 cm for pedals). When pushed, buttons and pedals interrupted a laser light beam delivered via fiber optic cable. The responses recorded from pedals and buttons were automatically labeled as ‘correct’ or ‘erroneous’ after each trial in accordance with the task rules (see below).

For the entire length of the experiment, the participants were comfortably seated in the MEG apparatus inside of an electromagnetically and acoustically shielded room (see below). Pseudowords were presented binaurally via plastic ear tubes in an interleaved quasi-random order, at 60 dB SPL. The experiment was implemented using the Presentation 14.4 software (Neurobehavioral systems, Inc., Albany, CA, USA).

The experiment comprised four blocks presented consequently in a fixed order for all participants: (1) passive listening before learning, (2) active learning, (3) active performance, and (4) passive listening after learning (Figure 1A). The whole experiment lasted approximately 2 hours. Here, we analyzed the data from two active blocks (results for the passive blocks were reported in Razorenova et al., 2020).

**Figure 1.**
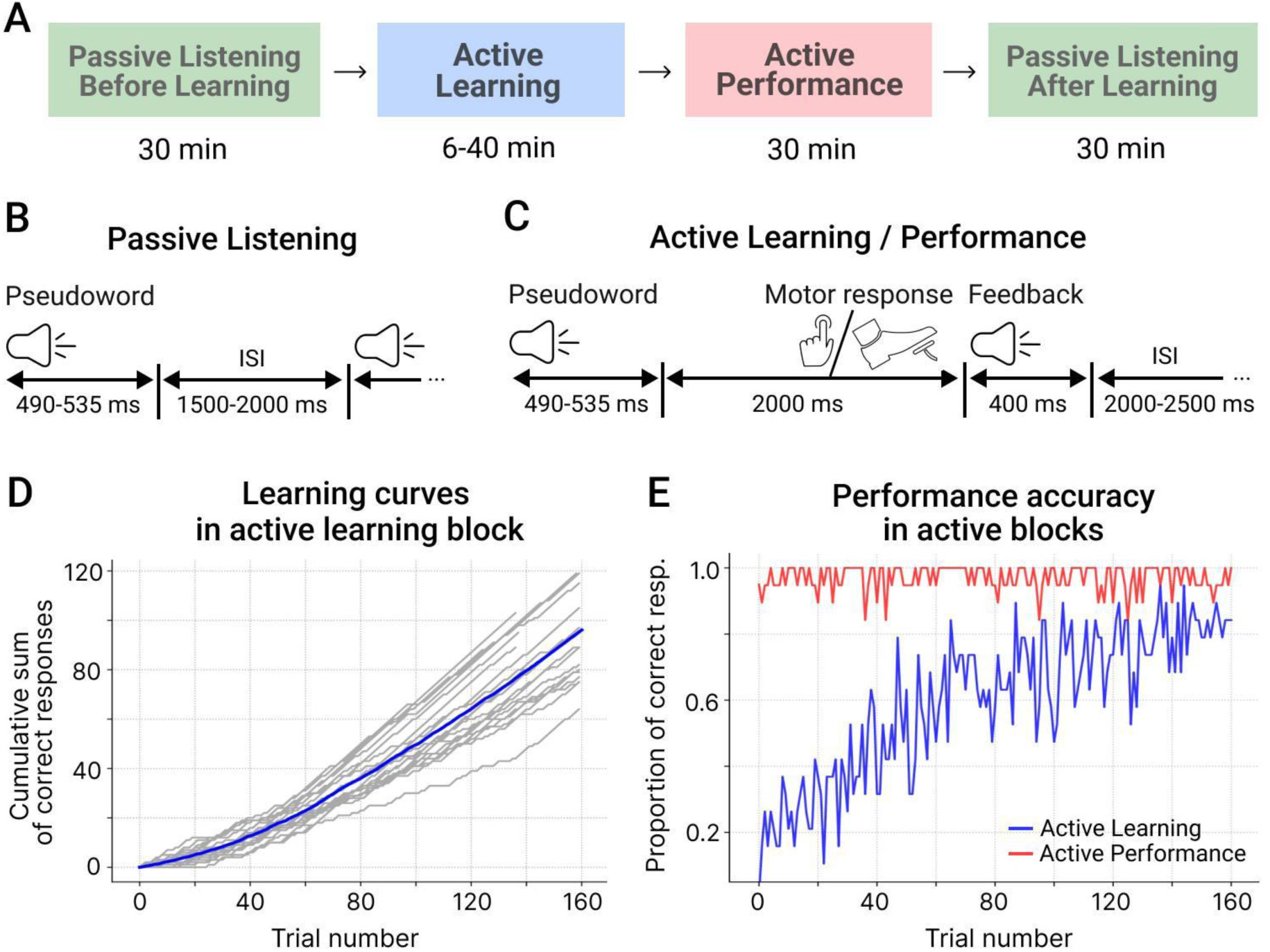
Experimental procedure and behavioral results. A. Sequence and approximate duration of experimental blocks. B. Trial structure in the passive listening block. C. Trial structure in the active learning and active performance blocks. D. Cumulative performance accuracy over the first 160 trials of the active learning block. The gray lines represent individual learning curves, the blue line - the grand average. E. The proportion of correct responses over the first 160 trials of the active learning (blue) and active performance (red) blocks.

During two passive listening blocks the participants were auditorily presented with the pseudowords in a pseudorandom interleaved order. An average interstimulus interval (ISI) was 1750 ms with a random jitter between 1500 and 2000 ms at one-millisecond steps (Figure 1B). Each passive block consisted of 400 stimuli (each of eight pseudowords was presented 50 times) and lasted around 30 min. To prevent the participants from paying attention to the auditory stimuli, they were presented with a silent movie of their choice projected on the screen positioned at eye-level.

Upon completion of the first passive block, the participants were instructed that during the following active blocks they had to deduce the unique associations between each of the presented eight pseudowords and movements of their own body parts. In order to do that, they were required either to respond to each pseudoword by using one of the four body extremities or to withhold motor response, and then listened to positive and negative feedback signals informing whether the action was correct or erroneous. The instruction did not contain any other clues: the behavioral procedure implied trying a variety of auditory cue-action associations and eventually settling down on those that led to positive reinforcement, thus complying with the requirements of operant learning (Neuringer, 2002).

During the active blocks, participants were instructed to keep their gaze at the fixation cross in the center of the presentation screen, with the purpose of minimizing artifacts caused by eye movements. The eight pseudowords were repeatedly presented in pseudorandom order. Each pseudoword was followed by a feedback signal. Positive feedback was given if a participant complied with the task rules, i.e., executed a correct movement to an APW stimulus or committed no response to an NPW stimulus (Table 1). Negative feedback followed three kinds of errors: (i) no response to an APW stimulus; (ii) a motor response to an APW stimulus performed with a wrong extremity; (iii) any response to an NPW stimulus. Both types of feedback were presented 2000 ms after the end of the pseudoword stimulus (Figure 1C). The average inter-stimulus interval (from the end of the feedback stimulus until the onset of the next pseudoword stimulus) was 2250 ms, randomly jittered between 2000 and 2500 ms at one-millisecond steps.

The number of stimuli in the active learning block varied across participants depending on their individual learning rate. The block ended if a participant reached the learning criterion or if 480 stimuli were presented in total, whichever came first. The learning criterion required that a participant made correct responses in at least four out of five consecutive presentations of each of the eight pseudowords. Correctness of the behavioral response as well as whether a participant met the learning criterion was automatically checked after each trial using custom-made scripts.

One participant failed to reach the learning criterion and thus went through all 480 trials in the learning block. Considering that his overall hit rate during the next active performance block was well within the range of performance of the other participants, we included this participant in all analyses. There was considerable variation between the participants in the duration of the active learning block: the number of trials required for a participant to reach the learning criterion ranged from 74 to 480, with the duration of the learning block varied from 6 to 40 min.

In the next active performance block (Figure 1A), the participants were asked to repeat the same procedure. The only difference between the two active blocks was that the active performance block included a fixed number of 320 trials and lasted approximately 30 min.

The participants were offered short breaks between the blocks (10 min between the active performance block and the second passive block and 3 min between other blocks), during which they rested while remaining seated in the MEG apparatus.

### MEG data acquisition

The experiment was conducted at the research facility “Center for Neurocognitive Research (MEG-Center)” of MSUPE. MEG was recorded in a magnetically shielded room (AK3b, Vacuumschmelze GmbH, Hanau, Germany) using a dc-SQUID Neuromag VectorView system (Elekta-Neuromag, Helsinki, Finland), which has 306 MEG channels (204 planar gradiometers and 102 magnetometers). The MEG signals were recorded with a band-pass filter 0.1–330 Hz, digitized at 1000 Hz, and stored for offline analysis.

Participants’ head shapes were measured using a 3Space Isotrack II System (Fastrak Polhemus, Colchester, VA, USA) by digitizing three anatomical landmark points (nasion, left, and right preauricular points) and additional randomly distributed 60-100 points on the scalp. Head position and orientation were continuously monitored during MEG recording by four Head Position Indicator coils. Two pairs of electrooculographic electrodes located above and below the left eye and at the outer canthi of both eyes were used to record vertical and horizontal eye movements.

Biological artifacts and other environmental magnetic sources originating outside the head were removed from MEG data using tSSS algorithm (Taulu, Simola, & Kajola, 2005) implemented in the MaxFilter program (Elekta Neuromag software). For further sensor-level analysis, MEG data were converted to a standard head position (x = 0 mm; y = 0 mm; z = 45 mm). Static bad channels were excluded from further processing. Correction of biological artifacts (caused by vertical eye movements, eye-blinks and heart-beats) was performed by means of the ICA method implemented in MNE-Python software (Gramfort et al., 2014). In order to exclude epochs with high muscle activity, we calculated the mean absolute signal values filtered above 60 Hz at each channel. The epochs during which maximal amplitudes at more than a quarter channels exceeded 9 standard deviations of the across-channel average were excluded from the analysis.

### Behavior analysis

The participants’ performance was evaluated as (i) a mean number of correct responses for four APW pseudowords, which were associated with movements (correct movement execution), for four NPW pseudowords, which were not (correct refraining from any movement), and the total number of correct APW and NPW trials, and (ii) a mean latency of the motor responses for APW pseudowords. This evaluation was performed separately for each of the two active blocks. The performance between two types of stimuli within each block, and between the blocks was compared using the paired t-test.

To characterize the learning curve, we used the cumulative record of correct responses as a function of trials (Gallistel, Fairhurst, & Balsam, 2004), which is calculated as a running sum of successive behavioral measurements. Changes in the slope of this curve correspond to changes in the level of performance (Figure 1D). Given individual variability in the learning curves, to visualize the general course of performance at the group level, we plotted a proportion of subjects, who gave a correct response at each consecutive trial, for each block separately (Figure 1E).

Considering that during the active performance block the participants reached a near-ceiling level of accuracy (Figure 1E), we resorted to an RT analysis in order to check whether learning progressed further during the advanced stage of the experiment. For this purpose, we applied linear mixed-effects models. We used an RT in each trial in the active performance block as a dependent variable with a corresponding trial number and a limb the participants responded with as fixed factors. Differences in RT between the participants were attributed to random effects. The full model was as follows:

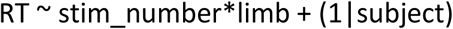

The linear mixed-effects models were built with the “lmer()” function available in the lme4 package for R (Bates, 2010).

### Epoch selection and MEG data preprocessing

To probe two different explanations of putative learning-induced contingency between auditory and motor activation during the time delay between a heard pseudoword and a performed motor response (or absence of a response), we evaluated both stimulus-locked and response-locked brain activity. In both analyses, we focused on the difference in MEG data between two conditions: the early and the advanced stages of auditory-motor association learning (ESL and ASL respectively), during which the participants gradually acquired the association rules between the pseudowords and the actions of body extremities (or alternatively with the absence of any actions). We adopted the following strategy for epoch selection.

For both stimulus-locked and response-locked analyses, we extracted the earliest trials from the active learning block (ESL condition) and the latest trials from the active performance block (ASL condition), and we did that extraction separately for two types of trials: with a motor response (APW) and without a motor response (NPW). We selected only the correct trials, while any incorrect trials were discarded. Since the participants used four extremities to make their responses, we counterbalanced the contribution of extremities into the dataset, thus avoiding any possible effects and interactions related to a particular extremity. For APW and NPW conditions separately, we extracted 28 correct trials (7 trials for each extremity) from each active block. If at least for one extremity the number of selected trials was less than 7, a respective participant was excluded from all further analyses. This procedure resulted in the exclusion of 9 participants, who completed the active learning block too quickly to allow extraction of a sufficient number of trials from the ESL condition, and the following analyses were done in 19 participants.

For both sensor-level and source-level analyses, the trials were epoched from −700 to 2500 ms relative to pseudoword onset for the stimulus-locked data; and from −1500 up to 1000 ms relative to the motor response onset for the response-locked data. Both stimulus-locked and response-locked event-related fields (ERFs) were baseline-corrected using 450 ms pre-stimulus window (from −500 ms to −50 ms before the stimulus onset); further details of the baselining procedure at sensor and source levels see below.

### Sensor-level analysis

First, we will describe the general framework of how we treated the sensor-level data in both stimulus- and response-locked analyses.

For the sensor-level analysis, MEG data were downsampled to 300 frames per second. At sensor-level, we used MEG signal from planar gradiometers which are known to be less sensitive to distant sources than magnetometers and, thus, provide clearer and easier-to-interpret picture (Garcés, López-Sanz, Maestú, & Pereda, 2017; Gross et al., 2013; Vrba & Robinson, 2001).

To examine and illustrate general ERF dynamics over time in a specific condition, we computed root mean square (RMS) over ERFs values across all 204 gradiometers at each time point, thus obtaining time courses of global field power (GFP). The baseline values were subtracted from GFP data. The resulting GFP time courses were compared between conditions of interest (Passive Listening vs ESL, ESL vs ASL) using the paired two-tailed t-test with FDR correction (Benjamini & Hochberg, 1995) for multiple comparisons for the number of time points.

For topographical analyses over the whole head, we computed RMS values of ERFs separately for each pair of gradiometers (hereafter referred to as combined gradiometers, or simply sensors). The resulting RMS values were averaged within the time windows of interest (described in details below) for each subject and experimental condition separately. Significance of the between-condition differences at each sensor was assessed using the paired two-tailed t-test with FDR correction for multiple comparisons for 102 channels.

#### Early stage of active learning vs. passive listening (task effect)

First, we examined whether the task load by itself led to changes in ERFs evoked by the auditory pseudowords. To this end, we compared ERFs evoked by the same stimuli in the passive listening and active learning blocks. Since there were no movements during the passive block, for this comparison we used only those stimuli from the active learning block that did not trigger movements - the NPW pseudowords. We extracted the first seven trials for each of the four NPW pseudowords from the passive condition and an equal number of trials from the active learning condition, and calculated GFP over the whole sensor array. The GFP time courses were compared between the passive and active conditions using the paired two-tailed t-test with FDR correction for multiple comparisons for 960 time points within the stimulus-related epoch from −700 to 2500 ms.

#### Stimulus-locked ESL vs. ASL contrast (learning effect)

To test for possible enhancement and prolongation of stimulus-locked activity during the post-stimulus period at the advanced stage of learning, we compared the evoked magnetic field response between ESL and ASL conditions for both types of stimuli. Based on the “retrospective” hypothesis, we expected stronger learning effect for newly learnt “action” APW pseudowords as compared to non-action related NPW stimuli. Indeed, since the latter type of stimuli did not require a motor response to the auditory cue, co-activation of auditory and motor preparation processes was not in play.

First, we collapsed all conditions, and quantified the timing of the peak value in the GFP time course of the magnetic brain activity evoked by the pseudowords. Next, since we were interested in detecting a putative enhancement of the ERFs in the proximity to the action required – i.e. greater signal amplitude during the descending slope after the peak – we selected a 400-ms time window starting at the ERF peak. The endpoint of this window (approximately 1250 ms after the stimulus onset) was selected so that it avoids overlapping in time with movement execution. We averaged GFP values over all time points within this 400-ms time window, for ESL and ASL conditions separately. These values were subjected to repeated-measures ANOVA, with two factors: learning stage (ESL and ASL) and stimulus type (APW and NPW). Post-hoc pairwise comparisons were made using Tukey’s honestly significant difference (HSD) test. To ensure that the APW-NPW difference in learning effect in the pre-selected post-peak time window was not inherited from the earlier processing differences, for each stimulus type we calculated time courses of the GFP signal in the ESL and ASL conditions and compared them at each time point (with FDR-correction for a number of time points).

Since both analyses performed showed no significant learning effect for NPW pseudowords, (see Results section), we further examined APW stimulus type only. To evaluate scalp topography of the ESL-ASL differences, we limited the pre-selected post-peak time interval to the shorter 200-ms period for which the significant learning effect was observed in the GFP time courses. For each sensor separately, we averaged RMS values within the chosen time window, and contrasted ESL and ASL conditions using paired t-test with FDR corrections for multiple comparisons over 102 combined gradiometers.

Additionally, we plotted topographical maps for successive 100-ms time windows, both in APW and NPW trials before and after learning, as well as differential response to learning for each trial type (APW ESL vs APW ASL; NPW ESL vs NPW ASL).

#### Response-locked ESL vs. ASL contrast

We tested whether the putative learning-induced increase in auditory cortex activation was present in response-locked ERFs. Only the APW trials, which involve motor responses, were considered. This analysis aligned MEG signals to the movement onset, while the information about the time point of word onset was jittered due to differences in reaction times.

First, for response-locked data, we compared GFP time courses for ESL and ASL conditions, using the two-tailed paired t-test, at each time point separately, with FDR correction for the number of timepoints in the whole epoch (750 time point within the response-related epoch from - 1500 to +1000 ms).

Next, in order to evaluate the topography of the learning effect, we chose the a priori window from −300 to −100 ms before the motor response, which corresponded to the movement preparation. This time window was well suited for between-condition comparison of the response preparation period: considering the reaction times in ESL and ASL conditions (1345 ± 105 ms and 1264 ± 117 мс, respectively), its lower boundary avoided the maximal auditory response, while the upper boundary spared an onset of EMG activity that occurred time-locked to a button press (Tandonnet, Burle, Vidal, & Hasbroucq, 2003). This time window roughly corresponded to the interval that we defined as the most significant for stimulus-locked data (see above). For each combined gradiometer, RMS values were averaged across the time window of interest in each condition separately and then subjected to the paired t-test with FDR correction for multiple comparisons for 102 combined gradiometers.

In addition, we plotted the topographical maps of successive 100-ms time windows over the second before the movement onset before and after learning, as well as the differential response between the ESL and ASL stages of learning (ASL minus ESL).

Any learning effects in the response-locked data could in fact be spurious, related to differences in response time between ESL and ASL conditions. Considering that the RTs were shorter in the ASL than in ESL condition, the increase in the response-locked activity in the ASL could be explained by the mere fact that response-locked ERF became closer to the stimulus. To control for the difference in the response latency, we repeated the analysis for the data matched for the condition-averaged RT (see Appendix 3).

#### Comparison between stimulus-locked and response-locked effects

At the next step, we compared the strength of the learning effects in the stimulus-locked and response-locked data. This comparison was crucial for differentiating between the “retrospective” and “prospective” explanations of the enhanced evoked response at the advanced stage of learning, since the effect we observed in the movement-related data, could be simply a carry-over effect of the power increase in the stimulus-locked ERFs. We expected that, if the increase in the auditory cortex activation was indeed related to the movement preparation, the learning effect should be stronger in the response-locked data compared with the stimulus-locked.

With this goal in mind, we directly compared stimulus-locked and response-locked ERFs before and after learning, within the two time windows of equal length of 200 ms: the window of the most significant ESL-ASL differences in the stimulus-locked response (see above) and the *a priori* chosen −300 - −100 ms window in the response-locked signal. For this purpose, we took only those combined gradiometers that were statistically significant in both analyses. The RMS values for these channels were averaged over respective 200-ms windows and subjected to repeated-measures ANOVA with two factors: learning stage (ESL and ASL) and ERF locking (stimulus-locked and response-locked). Post-hoc pairwise comparisons were made using the Tukey HSD test.

### Source-level analysis

#### MRI scanning, co-registration and source estimation

Participants underwent MRI scanning with a 1.5T Philips Intera system for further reconstruction of the cortical surface. Individual structural MRIs were used to construct single-layer boundary-element models of cortical gray matter with a watershed segmentation algorithm (FreeSurfer 4.3 software; Martinos Center for Biomedical Imaging, Charlestown, MA, USA). Individually recorded head shapes were then co-registered to this mesh using fiducial points and around 60 individually digitized scalp-surface points. A grid spacing of 5 mm was used for dipole placement, which yielded 10,242 vertices per hemisphere.

Both magnetometer and planar gradiometer data were used to compute sources. The evoked data were downsampled to 100 samples per second. A noise-covariance matrix was computed for each subject from the empty-room data recorded immediately before the experiment. The noise-covariance matrix and the forward operator were combined into a linear inverse operator using sLoreta algorithm implemented in the MNE software suite using default parameters. The source estimates were morphed onto the standard MNI brain using the surface-based normalization procedure implemented in MNE (Gramfort et al., 2014). The data were baseline-corrected at each vertex for the averaged current strength computed over the pre-stimulus −500 - −50 ms interval.

#### ESL vs. ASL contrast

We performed source reconstruction to allow for a more detailed description of the brain areas involved in the observed sensor-level effects of learning (ESL vs ASL differences) for both stimulus-locked and response-locked signals. Only trials with the motor responses were included in this analysis (i.e. APW trials).

First, for each time window of interest that was specified during the sensor-level analysis (for both stimulus-locked and response-locked activation, see above), we determined cortical areas involved in the neural response during ESL and ASL conditions separately. For each vertex, the cortical evoked current was averaged across the designated time windows at each condition and contrasted by means of the paired two-tailed t-test. To correct the results for multiple comparisons, we used the same FDR procedure as at the sensor level but across the entire set of vertices of the cortical surface instead of the MEG channels.

The cortical regions with the most reliable statistical effect were defined based on the probabilistic atlas of the human brain (Destrieux, Fischl, Dale, & Halgren, 2010) (see Results). Then, we extracted time courses of the source current from all vertices in each of the selected areas, under each condition separately. The resulting time series were averaged across the vertices for ESL and ASL conditions and, then, subjected to the point-by-point comparison by the paired two-tailed t-test, with FDR-correction for the number of time-points (250 time points corresponding to 2500-ms epoch).

#### Response-locked ERFs for movements made by each hand separately

For the analyses described above, the epochs were selected in such a way that hand and foot movements were pooled together in equal proportion, thus obscuring any movement-related somatotopic effects in the sensory-motor cortex. To uncover lateralized activity related to movement preparation, we conducted an additional analysis specifically focusing on trials involving hand movements, distinguishing between those performed with the left and right hand. We extracted all correct APW trials involving hand movements from the active learning and active performance blocks separately. To improve the signal-to-noise ratio, we used all relevant epochs from each block rather than limiting the analysis to seven epochs (24.2 ± 11.7 in the active learning block, 35.9 ± 4.5 in the active performance block, mean ± SD). Then, we reconstructed time courses of cortical response in four regions of interest (see Results for details). Point-by-point comparisons between ESL and ASL conditions were made as described above.

## Results

### Behavioral performance

All participants successfully acquired the rules linking the eight different pseudoword cues with the movements of specific body extremities – or with withholding a movement – through active operant learning. The analyses below are reported for those 19 participants who had a sufficient number of trials in the active learning block and were included in the main dataset (see Methods). Across the active learning block, the mean proportion of correct responses was 0.78 (SD = 0.6), being significantly different from that of chance (t(18)=14.98, p < 0.001). The correct response rate increased over the active learning block as learning proceeded. Figure 1D represents the cumulative record of correct responses as a function of trials in the active learning block in individual participants and at the group level. Inspection of the plots confirmed the general tendency for continuous improvement from an untrained, chance level of responding to a near perfect performance.

During the active performance block, all participants scored highly on the pseudoword-action association task (mean proportion of correct responses became as high as 0.97, SD = 0.04), indicating that they had successfully discovered and retained in memory the rules linking each of the eight auditory cues with the respective actions or non-action. As can be seen in Figure 2E, the participants maintained the near-ceiling performance level throughout the entire performance block, with the mean proportion of correct responses being well above 80%. There were no statistically significant differences in accuracy between APW and NPW trials both during the active learning block (70.8 ± 9.0 and 70.7 ± 6.7% for APW and NPW respectively, t(18)=0.06, p=0.95) and during the active performance block (96.0 ± 3.5 and 97.3 ± 4.3% for APW and NPW respectively, t(18)=1.59, p=0.13).

**Figure 2.**
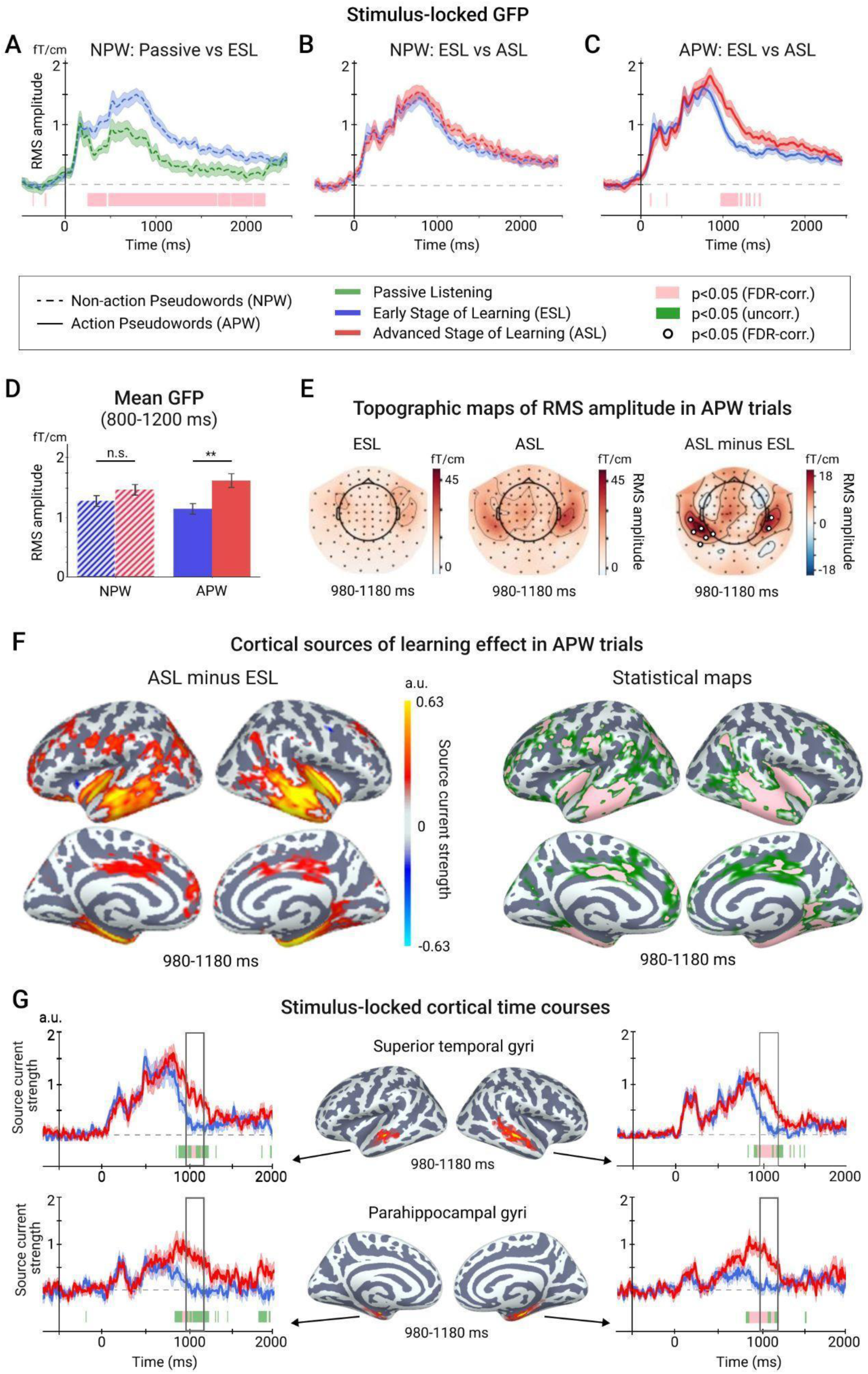
Learning effect in the stimulus-locked analysis. A. Time courses of global field power (GFP) in the first passive listening and early stage of learning (ESL) conditions in response to non-action related pseudowords (NPW). A vertical line corresponds to the stimulus onset. Here and hereafter, the shaded area around a time course represents the standard error of the mean (SEM). Colored horizontal bars under time courses represent time intervals with the significant between-condition differences (pink represents p<0.05, FDR-corr.; green - p<0.05, uncorr.). B. Time courses of GFP at early and advanced stages of learning (ESL and ASL) conditions in response to non-action related pseudowords (NPW). C. Time courses of GFP at ESL and ASL conditions in response to action related pseudowords (APW). D. Bar graphs represent mean values of GFP within the 800-1200 post-stimulus interval for NPW and APW stimuli in both conditions. Whiskers indicate 95% confidence intervals. Asterisks denote the level of significance: n.s. - non-significant, **p<0.01. E. Topographic maps of the mean RMS signal within the 980-1180 ms post-stimulus interval for APW stimuli during ESL and ASL conditions (left panels) and their difference (right panel). Here and hereafter, white circles indicate sensors with significant ESL-ASL differences (p<0.05, FDR-corr.). F. Cortical sources of ESL-ASL differences in response to APW stimuli in the 980-1180 ms post-stimulus interval. The left panel depicts the difference in amplitudes of source currents between ESL and ASL conditions (thresholded by p<0.05, uncorr.). The right panel demonstrates statistical maps of the between-condition differences: green color represents p<0.05, uncorr.; pink color - p<0.05, FDR-corr.). G. Time courses of source current strength from selected cortical regions. The cortical map in the middle displays the vertices with significant ESL-ASL differences in the 980–1190 ms interval within the selected region. The time courses represent an average across the significant vertices in the region.

The RTs in the APW trials significantly shortened from the active learning block to the active performance block (1387 ± 171 vs. 1285 ± 114; t(18)=3.46; p=0.003). As expected, the decision to execute the cued motor response became easier as learning proceeded. Notably, LMEM analysis revealed that there was a significant negative relation between a successive trial number and RT in that trial within the active performance block (F(1, 2910.31)=4.31, p=0.038) due to progressive shortening of motor response latency toward the end of this block. The factor of limb was also significant (F(3, 2910.02)=15.78, p<0.001), with leg movements being significantly slower than hand movements. However, there was no interaction between the two factors (F(3, 2910.21)=2.36, p=0.069) revealing that the tendency of shortening RTs throughout the active performance block was present for all extremities; this indicates that strengthening of auditory cue-motor association proceeded during the advanced learning stage at similar speed for all four movement types.

In addition, we compared mean RTs in the subset of trials chosen for the MEG analysis (see Methods for the details): the RTs taken from the beginning of the active learning block were significantly longer than the ones from end of the active performance block (1345 ± 105 ms vs 1264 ± 117 ms for APW ESL and APW ASL conditions respectively; t(18)=3.51, p=0.002).

### Stimulus-locked ERFs

#### Sensor-level analysis: Early stage of active learning vs. Passive listening (task effect)

As the first step, we contrasted the neural responses triggered by the pseudowords at the early stage of the learning to a control condition, when participants listened to the same auditory stimuli without any specific task in mind during the first passive block (Figure 1B). Since in the latter condition participants did not perform any movements, we included only NPW trials in this analysis. Point-by-point comparisons of the GFP time courses between the two conditions (Figure 2A) showed a highly significant task-related increase of the stimulus-locked response, which started at ∼200 ms after the stimulus onset and sustained until at least 1500 ms.

The straightforward explanation of greater strength and duration of the auditory neural response at the ESL compared with the passive condition might be the increased attentional and memory load during the task performance. If the role of attention is to facilitate the acquisition and memorization of cue-outcome associations (Sutherland & Mackintosh, 1971), it is plausible that the amount of attention paid to the auditory cues is greatest at the initial stage of learning. Following the same line of thinking, one could expect that the attentional boost effect on the neuromagnetic response should diminish with practice and task rule acquisition. In contrast, the associative learning hypotheses, as described in the Introduction, lead to the opposite prediction. Therefore, to separate the putative contribution of active learning from that of attention, we further compared stimulus-locked ERFs between the ESL and ASL conditions.

#### Sensor-level analysis: ESL vs. ASL (learning effect)

To assess whether auditory cue-evoked activity diminishes or increases over the course of learning, we employed the following strategy. First, we identified the time of the peak of auditory response by pooling all conditions and stimulus types together for this analysis (ESL and ASL, both APW and NPW), and then focused on the post-peak period—i.e., the downslope of the response. The auditory evoked activity peaked around 800 ms post-stimulus onset. Given the timing of the word-form uniqueness point at 360 ms, the peak latency of the auditory neural response corresponded to the semantic N400(m) component in EEG and MEG (Hultén, Schoffelen, Uddén, Lam, & Hagoort, 2019; Kutas & Federmeier, 2011; Maess, Herrmann, Hahne, Nakamura, & Friederici, 2006; O’Rourke & Holcomb, 2002).

Next, we chose the 400-ms post-peak window (i.e. from 800 ms to 1200 ms), so that the selected window would be shorter than the mean RT in all conditions and, thus, leave out the post-response activity. The GFP values were averaged across the selected time window, separately for each type of stimuli in each condition, and subjected to rmANOVA with two factors: learning stage and stimulus type. The main effect of the learning stage was significant (F(1,18)=12.66, p=0.002, partial η^2^=0.41), as well as its interaction with the stimulus type (F(1,18)=9.87, p=0.006, partial η^2^=0.35). Post-hoc comparisons using the Tukey HSD test revealed that the auditory response in the post-peak time window significantly increased after learning, but only for the APW stimuli which were followed by a motor action (p=0.005) (Figure 2D). The post-peak neural response to the NPW stimuli, to which the participants did not respond with their body extremities, was not reliably modulated by learning (p=0.51).

To determine whether the learning-induced increase in neural activity evoked by the APW stimuli occurred specifically in the post-peak period or across the entire course of ERF, we performed point-to-point comparison of the GFP time courses for NPW and APW stimuli separately. While the GFP evoked by the NPW pseudowords did not differ between the ESL and ASL conditions at any time point (p>0.05, FDR-corr.) (Figure 2B), the neural response to the APW pseudowords increased significantly at the advanced stage of learning, approximately from ∼980 to ∼1180 ms after the stimulus onset (p<0.05, FDR-corr.) (Figure 2C). Thus, the learning specifically enhanced the descending slope of the neuromagnetic response, with no effect observed at its peak or earlier — but only for the ‘action’ APW pseudowords associated with motor actions.

The topographical distribution of the learning-induced effect in the selected time window comprised two roughly symmetrical spatial clusters over left and right temporal regions (p<0.05, FDR-corr.) (Figure 2E). For illustrative purposes, we also depicted successive snapshots of the ERF scalp topography throughout the period of the post-stimulus response in the ESL and ASL conditions (Appendix 1). As one can see in the RMS time courses and series of topographical plots, the learning effect persisted for a substantial time between the stimulus presentation and the response onset.

#### Source-level analysis: ESL vs. ASL (learning effect)

To assess the neural sources of the learning-induced effects in the APW trials that we had statistically demonstrated in sensor space, we performed the source localization analysis. The cortical sources were localized for the time window between 980 and 1180 ms post-stimulus onset, during which we observed the greatest learning effect at sensor level (Figure 2F). Consistent with the scalp topographies in Figure 2E, learning predominantly modulated the activity in the lateral and medial regions of the left and right temporal lobes, particularly the superior and inferior temporal sulci (STS and ITS, respectively), the parahippocampal gyri (PHG) and posterior insula (p<0.05, FDR-corr.).

The time courses of the source current, averaged across the vertices with significant ESL-ASL difference in the STS and PHG regions (see Table 2 for MNI coordinates of the statistical maxima), were compared between the ESL and ASL conditions using paired t-tests (Figure 2G). The results aligned with those observed at the sensor level: the stimulus-related responses in the ESL and ASL conditions differed significantly between approximately 900 and 1200 ms post-stimulus onset, displaying similar dynamics bilaterally in both the STS and PHG regions (p < 0.05, FDR-corr.).

**Table 2.** Brain regions with the most reliable ESL-ASL differences in the stimulus- and response-locked data. MNI coordinates are given for the most significant vertices within the standard anatomical labels (Destrieux et al., 2010).

| Brain region | MNI coordinates<br>of the most significant vertex |  |
| --- | --- | --- |
|  | Stimulus-locked analysis | Response-locked analysis |
| Left parahippocampal gyrus | (-23.3, -12.7, -28.8) | (-20.5, -14.2, -26.8) |
| Right parahippocampal gyrus | (22.8, -20.3, -24.1) | (22.8, -20.3, -24.1) |
| Left superior temporal sulcus | (-51.9, -17.5, -8.0) | (-47.9, -24.9, -10.3) |
| Right Superior temporal sulcus | (44.6, -28.9, -6.6) | (46.4, -25.6, -10.6) |

In summary, both sensor- and source-level results indicated that learning of the stimulus-response associations slowed the decay of the neuromagnetic response during the interval between the stimulus and the cued action. This significant learning effect on stimulus-locked activation was observed only for auditory stimuli that prompted a specific body movement, not for those requiring no movements. The cortical areas most affected by learning during the APW trials were in the lateral and medial temporal regions of both hemispheres. The learning-related modulations emerged after the peak of the auditory response and persisted until roughly the end of the delay period between the stimulus and movement onset.

### Response-locked ERFs

#### Sensor-level analysis: ESL vs. ASL (learning effect)

The next step of the analysis was to examine the learning effect on the response-locked activity. As the correct NPW trials did not contain overt motor responses, this part of the analysis was conducted only for APW pseudowords.

On the response-locked APW epochs, the statistical t-test comparing point-by-point GFP amplitudes revealed a highly significant learning-related increase in the ASL compared with the ESL condition, which started about ∼600 ms before movement onset and lasted until the movement onset itself (p<0.05, FDR-corr.) (Figure 3A). Within this extended period of the significant learning effect, we selected a specific time window of interest—spanning from 300 to 100 ms before movement onset—to capture motor preparation. Figure 3B shows that the gradiometers exhibiting the most significant learning-induced changes were located bilaterally over the temporal lobes. Scalp topography maps at consecutive 100 ms time windows show that these statistical clusters persist throughout the entire pre-movement period (Appendix 2).

**Figure 3.**
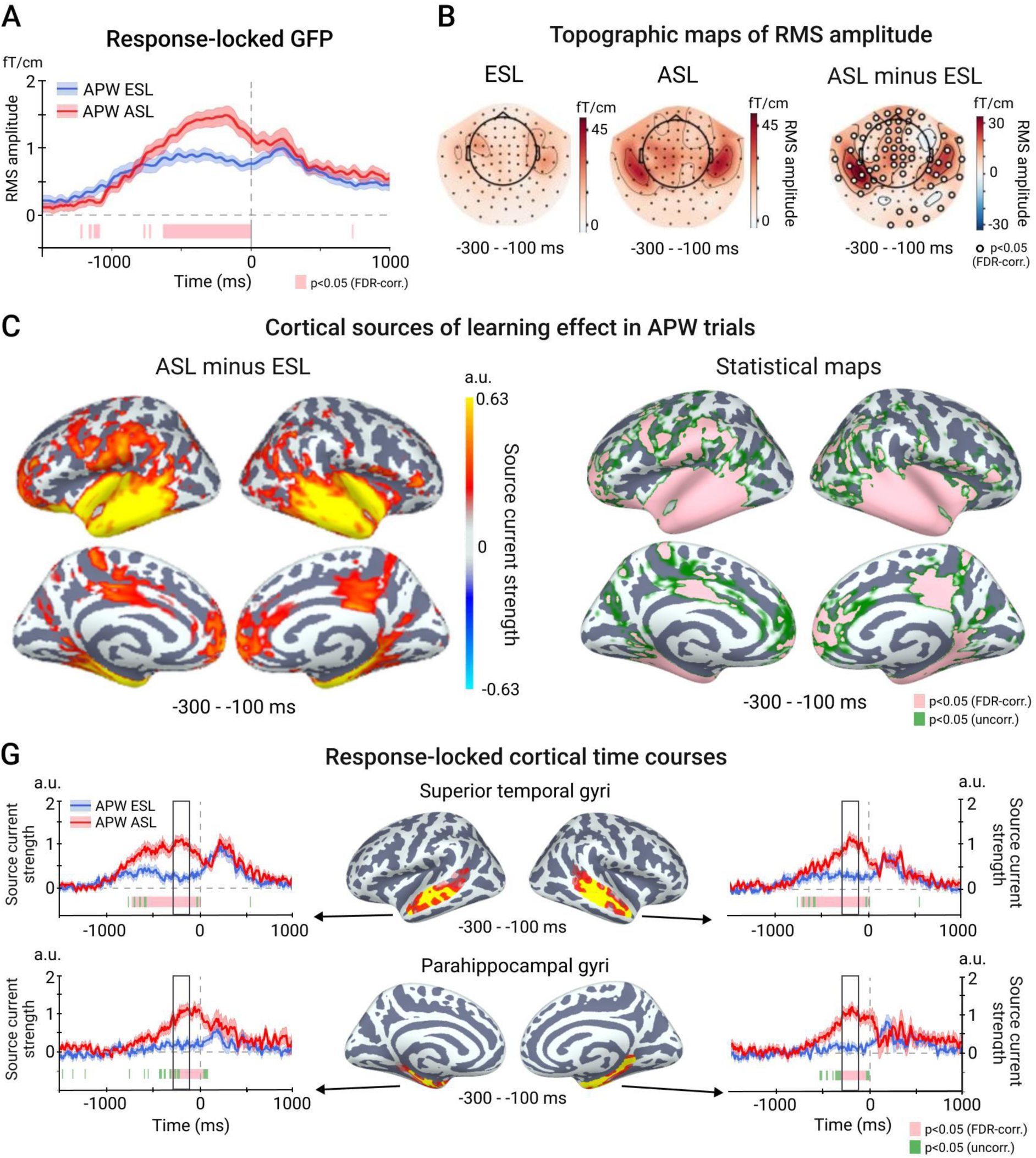
Learning effect in the response-locked analysis. A. Time courses of global field power (GFP) at early and advanced stages of learning (ESL and ASL) conditions during trials with action related pseudowords (APW). A vertical dashed line corresponds to a movement onset. B. Topographic maps of the mean RMS signal within the −300 - −100 ms pre-movement interval during ESL and ASL conditions (left panels) and their difference (right panel). C. Cortical sources of ESL-ASL differences in the −300 - −100 ms pre-movement interval. The left panel depicts the difference in amplitudes of source currents between ESL and ASL conditions (thresholded by p<0.05, uncorr.). The right panel demonstrates statistical maps of the between-condition differences: green color represents p<0.05, uncorr.; pink color - p<0.05, FDR-corr.). D. Response-locked time courses of source current strength from selected cortical regions. The cortical map in the middle displays the vertices with significant ESL-ASL differences in the −300 – −100 ms pre-movement interval within the selected region. The time courses represent an average across the significant vertices in the region.

The systematic reduction in RTs in the ASL condition, compared to the ESL condition, may have brought the pre-movement period closer in time to stimulus-driven processing, potentially causing spurious ESL-ASL differences in the movement-locked data. To ensure that the observed effect was not a technical result of RT difference, we selected subsets of trials in such a way that the group mean RTs did not differ between ESL and ASL conditions (mean RT ± SD: 1263 ± 119 ms vs 1256 ±123 ms; t(18)=1.18; p=0.25). The learning effect remained significant in the RT-matched data for approximately the same protracted period of time before the movement onset (approximately from −500 to 0 ms) as in the original data set (p<0.05, FDR-corr.) (Appendix 3).

#### Source-level analysis, ESL vs. ASL (learning effect)

The cortical areas underlying the learning effect on movement-locked ERFs in the time window of interest (−300 - −100 ms) are shown in Figure 3C. The set of cortical regions underlying the greater movement-locked ERF in ASL versus ESL conditions closely resembled that observed for the stimulus-locked response, though it was more extensive. Specifically, learning enhanced the movement-locked activation in widespread areas of the temporal lobe (including the STS, ITS, and PHG), the frontal pole, the posterior region of the mid-cingulate cortex, and the left inferior parietal cortex. The reconstructed time courses of the cortical activity from the STS and PHG areas (Figure 3D; see Table 2 for MNI coordinates of the most significant vertices) revealed that the significant learning-related differential activation (p<0.05, FDR-corr.) started at the left STS ∼700 ms before the movement onset, then, at around 450 ms, it became apparent in the right STS and in the PHG bilaterally. Once emerged, the learning effects persisted until the onset of the movement response, peaking approximately 200 ms before the movement execution.

Overall, our movement-locked data showed that the learning-induced increase in temporal lobe activity was present not only in response to the auditory stimulus but also in anticipation of the upcoming motor response. To distinguish between the “retrospective” and “prospective” accounts of this increase, a direct comparison of stimulus- and response-locked activity was necessary.

#### Learning effect in response-locked vs. stimulus-locked data

To contrast the learning-induced effects in the stimulus- and response-locked neural responses on a comparable time scale, we used time windows of equal length (200 ms) for both types of time-locking. We compared the response-locked data averaged across the pre-movement period of −300 - −100 before the response onset with the stimulus-locked data averaged across the interval of the most reliable ESL-ASL difference in the stimulus-locked analysis (980 - 1180 ms after the stimulus onset). Considering the mean RT of ∼1300 ms, two temporal intervals roughly overlapped. The spatial clusters of gradiometers showing significant effects in the ASL vs. ESL contrast (p < 0.05, FDR-corr.) had similar maxima for both types of locking (Figure 4A, C). However, the learning effect in the response-locked RMS was notably more extensive, involving gradiometers covering bilateral temporal and midline frontal regions.

**Figure 4.**
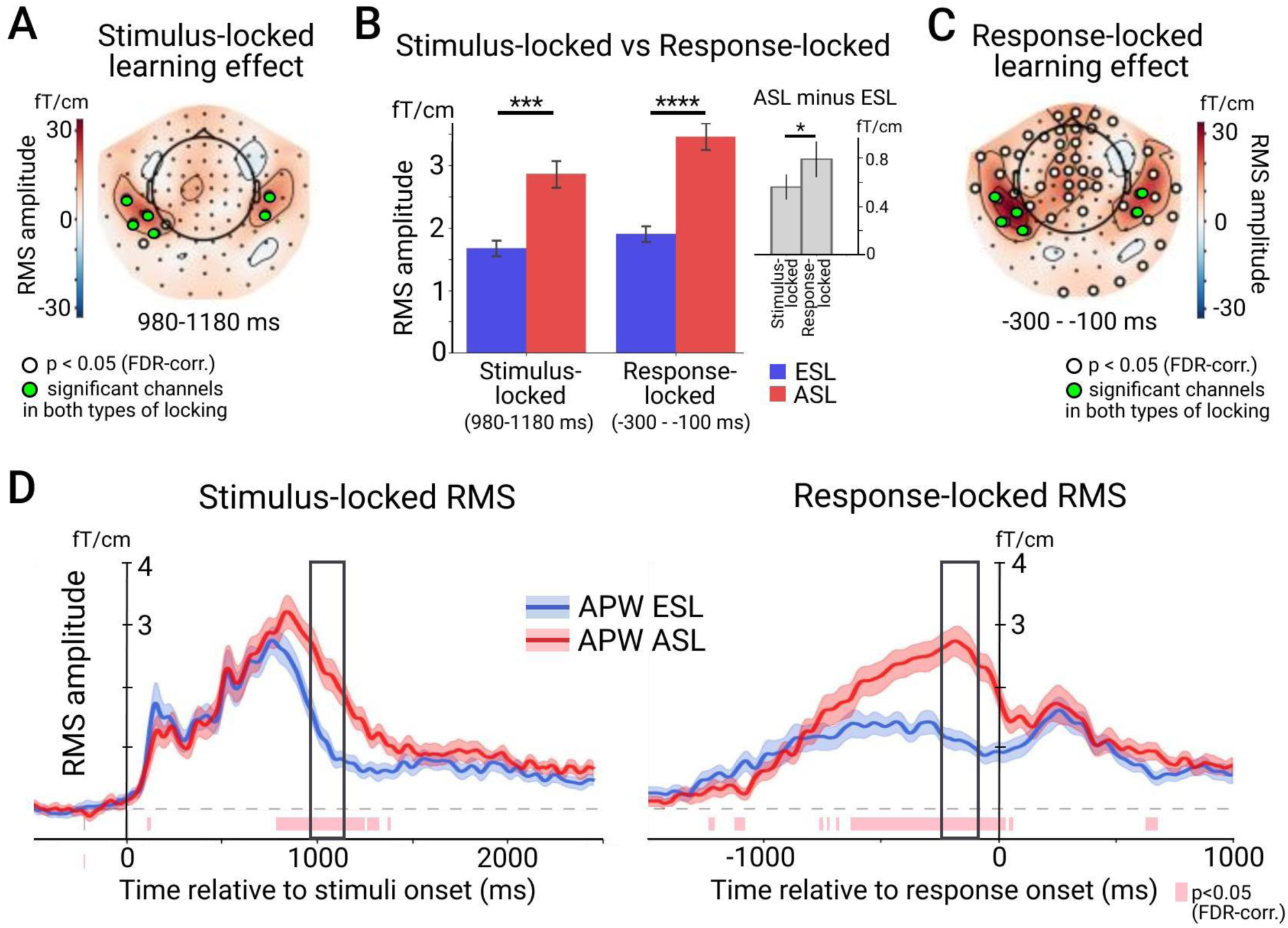
Comparison of the strength of the learning effect between the stimulus-locked vs. response-locked data. A. The topographical map of differential RMS amplitude between ESL and ASL in the stimulus-locked data during the time interval of interest. Here and elsewhere, white circles indicate channels with significant ESL-ASL differences (p<0.05, FDR-corr.), green circles mark the subset of channels which were significant in both analyses. B. Bar graphs represent mean values of RMS over the selected subset of channels at time of interest during ESL and ASL conditions in stimulus- and response-locked analyses. On the insert, a bar graph shows the difference in RMS amplitudes between ESL and ASL conditions in the stimulus- and response locked data. Whiskers indicate 95% confidence intervals. Asterisks denote the level of significance: *<0.05, **p<0.01, ***p<0.001, ****p<0.0001. C. The topographical map of differential RMS amplitude between ESL and ASL in the response-locked data during the time interval of interest. D. Time courses of RMS calculated over the subset of channels which displayed significant ESL-ASL differences in both stimulus- and response-locked analyses. The time course on the left is aligned to the stimulus onset, the time course on the right - to the onset of the motor response. The black rectangles indicate the time intervals of interest.

To further compare the stimulus-locked and response-locked data, we focused on those sensors that were significant in both analyses; they overlay the left and right temporal cortex. Point-by-point comparisons of the RMS time courses, representing the average across this subset of temporal gradiometers, showed a significant increase in the neural response in the ASL condition compared to the ESL condition for both stimulus- and movement-locked activity within the protracted interval preceding movement onset (Figure 4D).

Such activation of the auditory-related temporal cortex prior to the movement onset, might result from the sustained stimulus-locked activation “leaking” into the movement-locked neural responses. To check for such possibility, we directly compared the strength of the learning effects between the stimulus-locked and movement-locked ERFs using the same subset of “temporal” sensors. The RMS values were averaged across the temporal sensors and 200-ms windows of interest in the stimulus-locked and movement-locked ERFs separately and subjected to ANOVA with the learning stage (ESL vs. ASL) and the type of locking (stimulus-vs. movement-locked ERFs) as repeated measure factors. Both main factors were significant (learning stage: F(1,18)=31.89, p<0.001, partial η^2^=0.64; locking: F(1,18)=16.12, p=0.001, partial η^2^=0.47), as well as their interaction (F(1,18)=7.06, p=0.016, partial η^2^=0.28). The learning-related amplitude increase was significantly greater in the movement-locked compared with the stimulus-locked activity (t(18)=-2.72, p=0.014) (Figure 4B). The larger learning effect in the response-locked ERFs compared with the stimulus-locked ERFs in the temporal sensors refuted the possibility that the response-locked effect solely resulted from a spillover of the late parts of the stimulus-locked activity. To the contrary, the data do not exclude the possibility of leakage in the reverse direction—from the response-locked to the stimulus-locked activity. These findings suggested the presence of a distinct mechanism for the auditory cortex reactivation related to the movement preparation, consistent with the “prospective” hypothesis.

#### Learning effect for hand movements

A key prediction of the “prospective” hypothesis is that an acquired association between an auditory cue and the movement of a specific body part should lead to a temporal coincidence of motor and auditory activation within the involved cortical areas. Yet, in the analyses reported above, the movement-locked signal was averaged across four different body parts, and a respective somatotopic motor activation was blurred. Therefore, to examine the predicted co-occurrence in activation of auditory cortex and motor representation of the specific body part triggered by the readiness to initiate its movement, we focused on a well-studied MEG/EEG component called motor readiness potential, or motor readiness field (MRF) in MEG studies. The MRF is a contralaterally dominant slow field shift preceding unilateral movements that is considered as the MEG signature of motor preparation (Cheyne, Bakhtazad, & Gaetz, 2006; Kristeva-Feige et al., 1997; Schurger, Hu, Pak, & Roskies, 2021). Cortical sources of the MRF are localized to the hand area in the primary motor and premotor cortex predominantly in the contralateral hemisphere (Cheyne et al., 2006). Based on the notion of lateralized MRF as a strictly motor preparation component, we wanted to ensure that this MEG hallmark of motor activation was overlapping in time with the augmented auditory cortex activation preceding the hand movements.

To this end, we extracted movement-locked signals for the right and left hand movements in both ESL and ASL conditions, and reconstructed time courses of average source current for the four cortical regions selected from the Destrieux atlas (Destrieux et al., 2010). The first two regions comprised the superior part of the precentral sulcus from left and right hemispheres (Figure 5). These areas contain lateral premotor hand area (Yousry et al., 1997), known to contribute to generation of the hand-related MRF (Cheyne et al., 2006). The two other cortical regions included the whole STS label from the Dextraux atlas, chosen because of their association with lexical-phonetic aspects of speech processing, as demonstrated in fMRI and MEG studies (Poeppel, Idsardi, & van Wassenhove, 2007; Turkeltaub & Branch Coslett, 2010; Zhang et al., 2011). As MEG has low sensitivity to radially oriented sources in the cortical gyri (Hämäläinen, Hari, Ilmoniemi, Knuutila, & Lounasmaa, 1993), the regions of interest were limited by the sulci surface.

**Figure 5.**
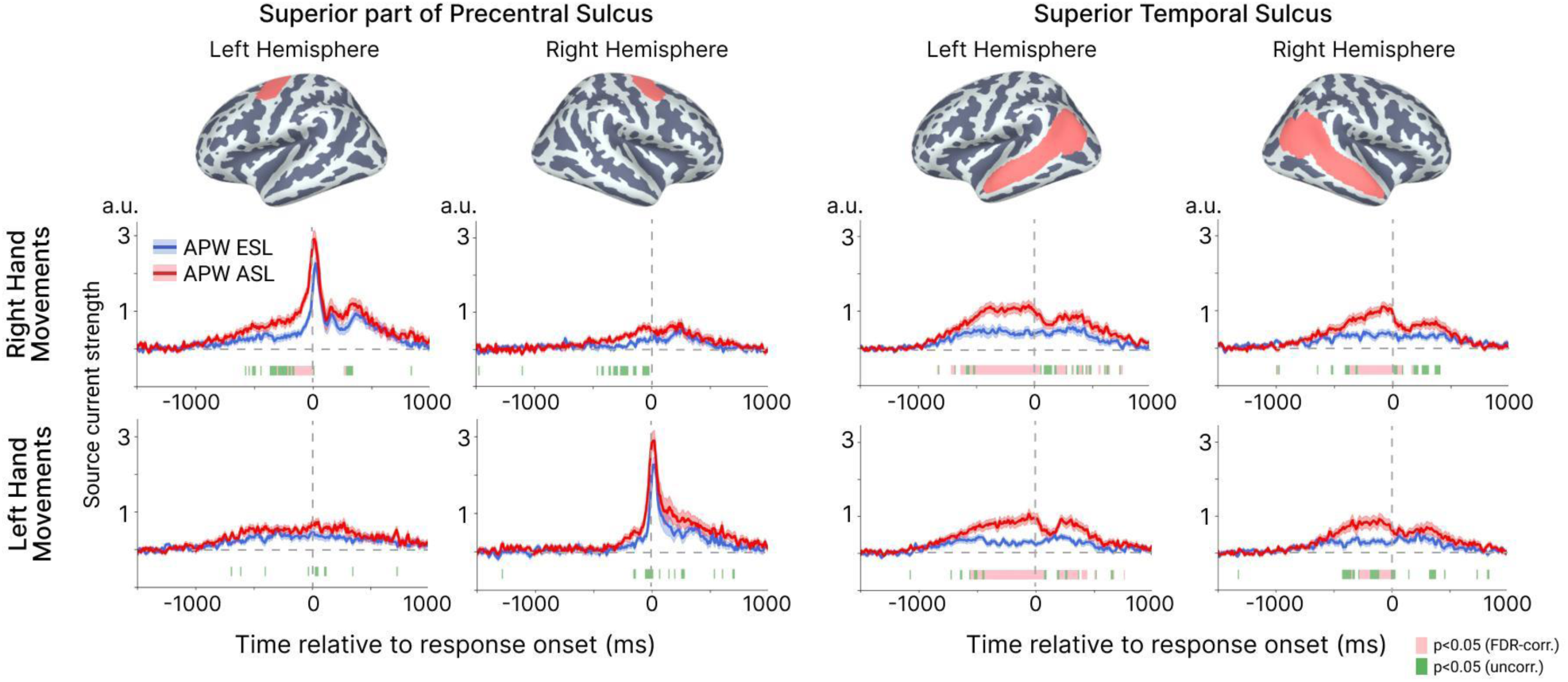
Learning effect during movements of the right and left hands. Each row represents response-locked time courses of source current strength from the superior part of precentral sulcus and superior temporal sulcus during right hand (upper row) and left hand movements (bottom row). The corresponding cortical regions are depicted on the lateral surface of both hemispheres. The time courses represent the average across the vertices in each region. A vertical dashed line indicates the movement onset. Colored horizontal bars under time courses represent time intervals with the significant between-condition differences (pink represents p<0.05, FDR-corr.; green – p<0.05, uncorr.).

Figure 5 shows the resulting time courses of cortical activity in the selected cortical regions during the right and left hand movements under the ESL and ASL conditions. Consistent with previous literature (Cheyne et al., 2006; Praamstra, Schmitz, Freund, & Schnitzler, 1999), we observed a bilateral slow amplitude shift (motor readiness activity) starting approximately 500 ms before movement. This shift was more pronounced for right-hand movements and eventually developed into a sharp increase in activity in the contralateral premotor hand area, peaking shortly after movement onset. The learning-induced STS activation also began around 500 ms before movement and persisted until the motor response was executed, with its onset and duration largely overlapping with that in the contralateral pre-motor hand area.

Thus, the enhancing effect of associative learning on the movement-locked bilateral activation in the auditory cortex coincides temporally with the motor preparation processes specific to the referential movement.

## Discussion

The current study sought to explore the putative augmenting effect of associative learning on the activation of the temporal cortex during the time interval between the auditory pseudoword and the motor response. We indeed found such an increase both in stimulus-locked and movement-locked neural responses mostly in the temporal cortices (including speech-related areas), although this increase was significantly more prominent in the response-locked data compared with the stimulus-locked data.

Comparing the neuromagnetic response at the early stage of learning to passive listening of the same pseudowords, we found that the requirement to memorize pseudoword-action associations enhanced and prolonged the stimulus-locked brain activity that peaked around 800 ms after pseudoword onset. In our study, the word-form uniqueness point of the spoken pseudowords corresponded to 360 ms after their onset, being the first moment when acoustic information required for pseudoword identification became available. Given this, the latency of the peak activity we observed agrees with N400-like activity described for spoken word/pseudoword processing, which typically peaks at around 400 ms after the recognition point (O’Rourke and Holcomb, 2002). The observed augmentation of the stimulus-locked activity under the learning task is in accord with previous studies that showed enhancement of the N400 component when attention is drawn to verbal stimuli, as opposed to its unattended processing (Erlbeck, Kübler, Kotchoubey, & Veser, 2014; McCarthy & Nobre, 1993). Furthermore, the N400 component with an unusually long descending slope has been observed in lexical decision tasks where, as in our study, a pseudoword representation was actively maintained in short-term memory to guide selection of an appropriate overt verbal response (O’Rourke and Holcomb, 2002). Therefore, the lengthened N400m in our study likely results from the increased demands on attention and memory during the early learning stage, compared to the passive listening.

This interpretation, however, does not explain the further increase in the stimulus-evoked neural responses observed over the course of learning. Both focused attention and memory retention were likely required throughout the entire period of mastering the pseudoword-action associations. Yet, as learning progresses and stimulus-to-movement matching becomes more automatic, the load on memory and attention typically decreases (Logan, 1979). This decrease should, in turn, reduce neural activation related to these cognitive processes. Therefore, it is unlikely that these general-domain processes alone account for the slowed decay of sustained stimulus-locked neural activation we observed at the advanced stage of learning—when errors had nearly disappeared and reaction times had significantly decreased, clearly indicating a high degree of automatization of the cued responses (Figure 1D, E).

In contrast, this finding appears to agree with the “retrospective” associative hypothesis, which posits that during learning, the activation of an input representation should be extended to overlap temporally with programming of the motor output. Accordingly, we observed that the enhanced N400m response to pseudowords was sustained until the onset of the corresponding movement, particularly at the late stage of learning when the unique associations between each pseudoword and its corresponding body part movement had been reliably established (Figure 2C). In other words, this sustained activation emerged only when confidence in the accuracy of the association was high, and the auditory pseudoword cue unambiguously predicted the execution of the corresponding action. Moreover, since no differences were observed between the early and late stages of learning for non-action-related pseudowords (Figure 2B), we infer that the learning-induced changes in the N400m wave were specific to pseudowords requiring the selection, preparation, and execution of movements.

However, the learning-induced enhancement of temporal cortex activation was also evident in the movement-locked data, which could support the “prospective” hypothesis. As with the stimulus-locked data, this enhancement became prominent only at the advanced learning stage, when movements were strictly determined by the preceding pseudoword, spanning a similar time interval and exhibiting a similar cortical topography in both data types (cf. Figure 4A, C). Notably, the learning effect was more pronounced in the movement-locked response than in the stimulus-locked response (Figure 4B). These suggest that the stronger movement-locked activation, induced by associative learning, might leak into the late stages of stimulus-triggered activity, potentially inflating or even creating the observed learning effect in the stimulus-locked response. While this does not entirely rule out the “retrospective” hypothesis, it lends stronger support to the “prospective” explanation.

The learning-induced increase in the movement-locked neural activity before the movement onset originated predominantly from medial and lateral temporal cortices in both hemispheres (Figure 3C). Parahippocampal gyrus on the medial surface with direct connections to the hippocampus is considered as a part of hippocampal memory circuits (Rolls, Deco, Huang, & Feng, 2022) and has been previously implicated in acquiring associations between diverse elements (Aminoff, Kveraga, & Bar, 2013). It is generally agreed that the hippocampal system is a convergence zone that receives signals from all the modalities and supports binding diverse representations into associative memories (Battaglia, Benchenane, Sirota, Pennartz, & Wiener, 2011; Wang et al., 2014). One of the possible approaches sees the hippocampus as an “indexing system” which points to the distributed neocortical regions that store memorized representations (Teyler & DiScenna, 1985; Teyler & Rudy, 2007). We speculate that the enhancement of parahippocampal gyrus activity at the advanced stage of learning reflects the process by which pairs of simultaneously activated representations of the acoustic pseudowords and corresponding motor programs become jointly indexed and associated.

On the lateral surface of the temporal lobe, the regions contributing to the learning effect on movement-evoked neural response to action pseudowords included the superior and inferior temporal sulci, known to be involved in the phonological and lexico-semantic processing of real words (Jefferies, 2013; Kutas & Federmeier, 2011; Silva Pereira et al., 2017; Zhang et al., 2011). This involvement indicates that, as pseudowords temporarily took on the meaning of action words, speech-processing resources were actively engaged during movement preparation, up until the moment just before movement onset.

Importantly, when the right and left hand movements were considered separately, we observed that movement-locked augmentation of the temporal cortex activity co-occurred with the lateralized motor readiness component in the contralateral pre-motor hand area – a signature of movement programming (Praamstra et al., 1999; Schurger et al., 2021) (Figure 5). This finding further supports the “prospective” association hypothesis that predicted a coincidence in time of the re-activation of the memorized input representation and preparation of the motor response.

The protracted, gradually increasing activation that we observed in the temporal cortex before an upcoming movement resembles the slow rising waves that occur in anticipation of predictable auditory stimuli (Bianco et al., 2020; C H M Brunia & van Boxtel, 2004; Ohgami, Kotani, Hiraku, Aihara, & Ishii, 2004). Such sustained “anticipatory” deflections belong to a broad class of slow event-related preparatory potentials (including stimulus preceding negativity, contingent negative variation, motor readiness potential and others), which are typically observed prior to predictable events. They are thought to reflect pre-activation of a specific representation which serves as a forward model of expected input or output (Brunia, van Boxtel, & Böcker, 2011; León-Cabrera, Hjortdal, Berthelsen, Rodríguez-Fornells, & Roll, 2024). In our task, sustained activation of the speech-related areas of the temporal cortex was triggered by the anticipation of movement associated with a specific auditory pseudoword, rather than by the auditory pseudoword itself. This suggests that at an advanced stage of learning, once the internal model linking acoustic pseudowords to a certain action is established, both the representation of word-form and its motor counterpart are activated during movement preparation. Our finding suggests the intriguing possibility that sustained anticipatory pre-activation may not be limited to cortical areas of the same modality as the predicted event but may also involve cortices that encode associations related to that event.

In this respect, the learning-induced re-activation of the auditory cortex before the upcoming movement reported here may be akin to a phenomenon of auditory-motor resonance – the bidirectional auditory-to-motor mapping that results in activation of motor cortical regions during listening to sounds produced by actions (Kaplan & Iacoboni, 2007) and in activation of auditory cortices while performing actions (i.e. as playing a familiar melody on a mute piano keyboard (Bangert et al., 2006)) (for extensive reviews, see Aglioti & Pazzaglia, 2010; Burgess, Lum, Hohwy, & Enticott, 2017). It has been shown that even short-term learning can lead to action-triggered auditory neural activity. For example, after short trumpet rehearsal, the mute trumpet-based task involving only finger movements increased activation of the primary auditory cortex in non-musicians (Gebel, Braun, Kaza, Altenmüller, & Lotze, 2013). A novel finding in the current study, however, was that as a result of newly acquired associations between sound and action, the auditory activity occurred in anticipation of the participant’s own movement, rather than being triggered by the movement onset. This finding demonstrates that during the planning of upcoming actions in an active operant conditioning procedure, the brain reactivates representations of auditory cues in the auditory cortex, along with their associated motor counterparts.

Taken together, our findings show that, as a result of the acquisition of one-to-one associations between auditory pseudowords and specific actions through trial-and-error search, the sustained temporal cortex activation is initially triggered by the presented pseudoword, and then re-occurs synchronously with initiation and preparation of the correct movement. This provides the first evidence that the adult brain, in order to form a novel association, relies on a “prospective” mechanism, which enables it to fill even a prolonged temporal gap between an auditory word and associated action. Such a reactivation may allow binding through Hebbian plasticity, i.e., the long-lasting synaptic changes in the temporally correlated neural assemblies.

## Author contribution

V.D. Tretyakova: Formal analysis; data curation, software; visualization; writing – original draft. A.A. Pavlova: Formal analysis, visualization; writing – original draft. V.V. Arapov: Formal analysis; software; visualization. A.M. Rytikova: data collection, project administration. A.U. Nikolaeva: investigation, software, data collection. A.O. Prokofyev: Investigation; software. B.V. Chernyshev: Conceptualization; methodology; supervision; funding acquisition. T.A. Stroganova: Conceptualization; methodology; data analysis, supervision; writing – original draft. All authors participated in the reviewing and editing of the final version of the manuscript.

## Acknowledgments

We are grateful to Anna Butorina for the help with the data analysis.

## Diversity in Citation Practices

The authors of this paper report its proportions of citations by gender category to be: M/M = 0.66; W/M = 0.191; M/W = 0.085; W/W = 0.064.

## Appendix 1. Topographic maps of stimulus-locked RMS signal

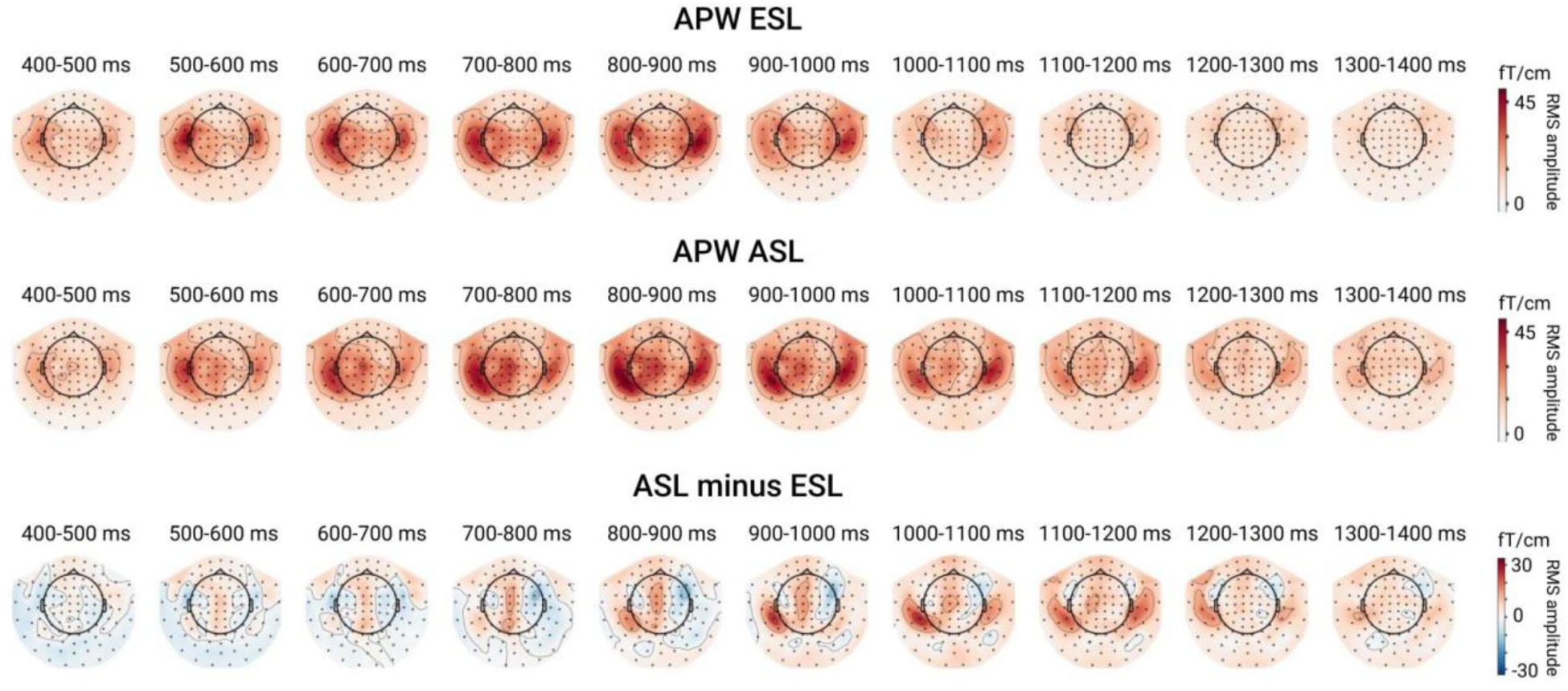

Topographic maps of RMS signal at consecutive 100 ms time windows relative to the stimulus onset (t=0 ms). The upper row represents ERF topography in response to action-related pseudowords (APW) at the early stage of learning (ESL). The middle row represents the response to APW at the advanced stage of learning (ASL). The bottom row - the difference between ESL and ASL. Note the different scale for the differential activity.

## Appendix 2. Topographic maps of response-locked RMS signal

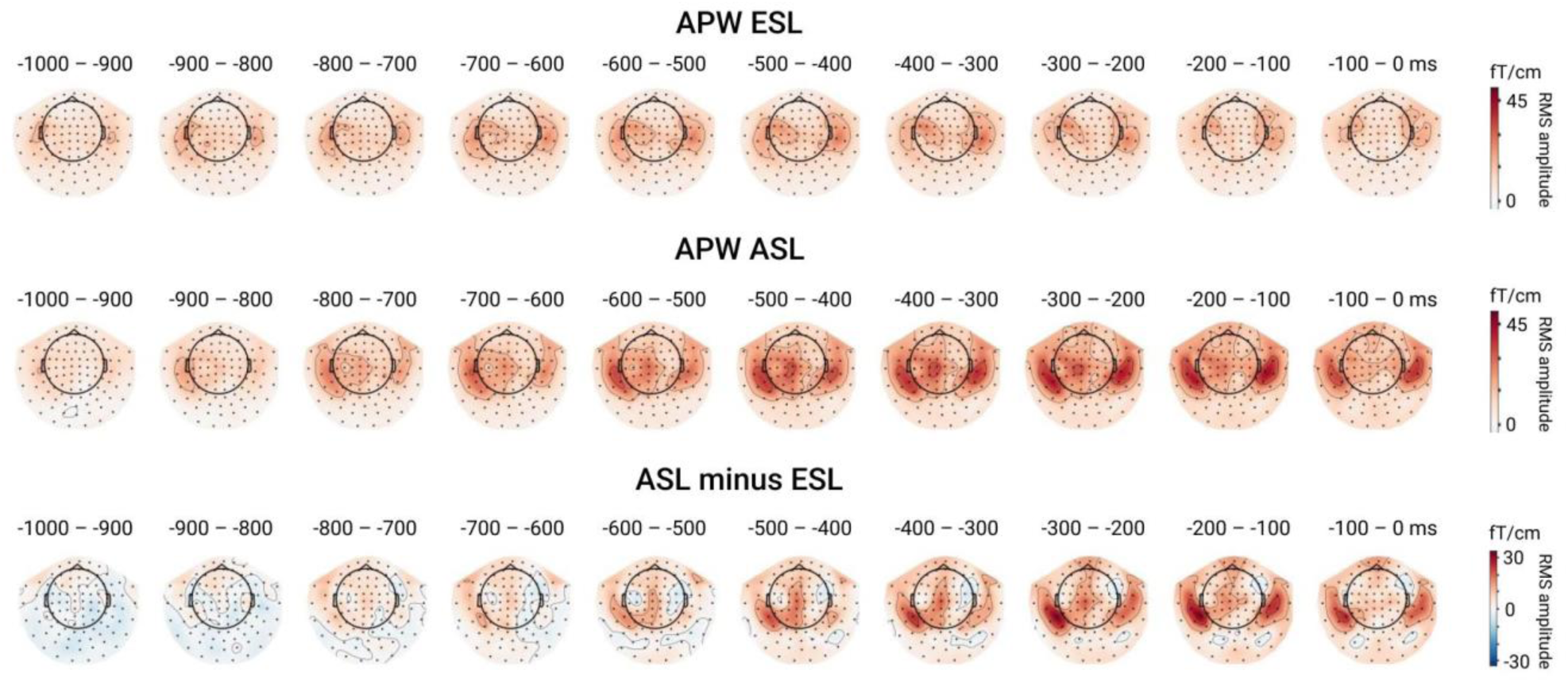

Topographic maps of RMS signal at consecutive 100 ms time windows relative to the movement onset (t=0 ms). The upper row represents ERF topography in response to action-related pseudowords (APW) at the early stage of learning (ESL). The middle row represents the response to APW at the advanced stage of learning (ASL). The bottom row - the difference between ESL and ASL. Note the different scale for the differential activity.

## Appendix 3. Response-locked ESL vs. ASL contrast: analysis on trials matched for the RT

The effects that we analyzed in response-locked data and interpreted as the effect of learning during motor preparation could in fact be spurious, related to differences in response time between ESL and ASL conditions. As long as the RTs were shorter in the ASL than in the ESL condition, the response-locked activity of interest could become closer to the stimulus and thus explain the observed increase in the ASL condition. To control for the difference in the response latency, we repeated the GFP analysis on the subset of trials that did not differ in the average RT between ESL and ASL conditions (i.e. only relatively faster responses from the ESL condition and relatively slower responses from the ASL).

To achieve this, we used the following response time matching procedure in each subject. For each trial from the ESL condition, we attempted to select a matching trial from the ASL: a matched pair was assigned on condition that the difference in reaction times within a pair of selected trials was less than 150 ms. The trials were picked one by one in the order in which they were presented to the subject. At each step, matched pairs of trials were removed from the further search. After this procedure was completed, only matched trials were included in the further analysis, while unmatched trials were discarded. This procedure resulted in average 21.16 ± 2.58 trials per subject. The RTs in those selected trials were averaged in the ESL and ASL conditions and, then, subjected to two-tailed paired t-test. After assuring that the RTs in the subset of trials did not differ between ESL and ASL (mean RT ± SD: 1263 ± 119 ms vs. 1256 ±123 ms; t(18)=1.18; p=0.25), we repeated comparisons of time courses of GFP as described in Methods. As can be seen in Figure below, the learning effect remained significant in the RT-matched subset of trials (both in stimulus- and response-locked analyses), supporting the notion that the observed increase in the movement-locked ERF was induced by learning rather than being caused by disparity in RTs.

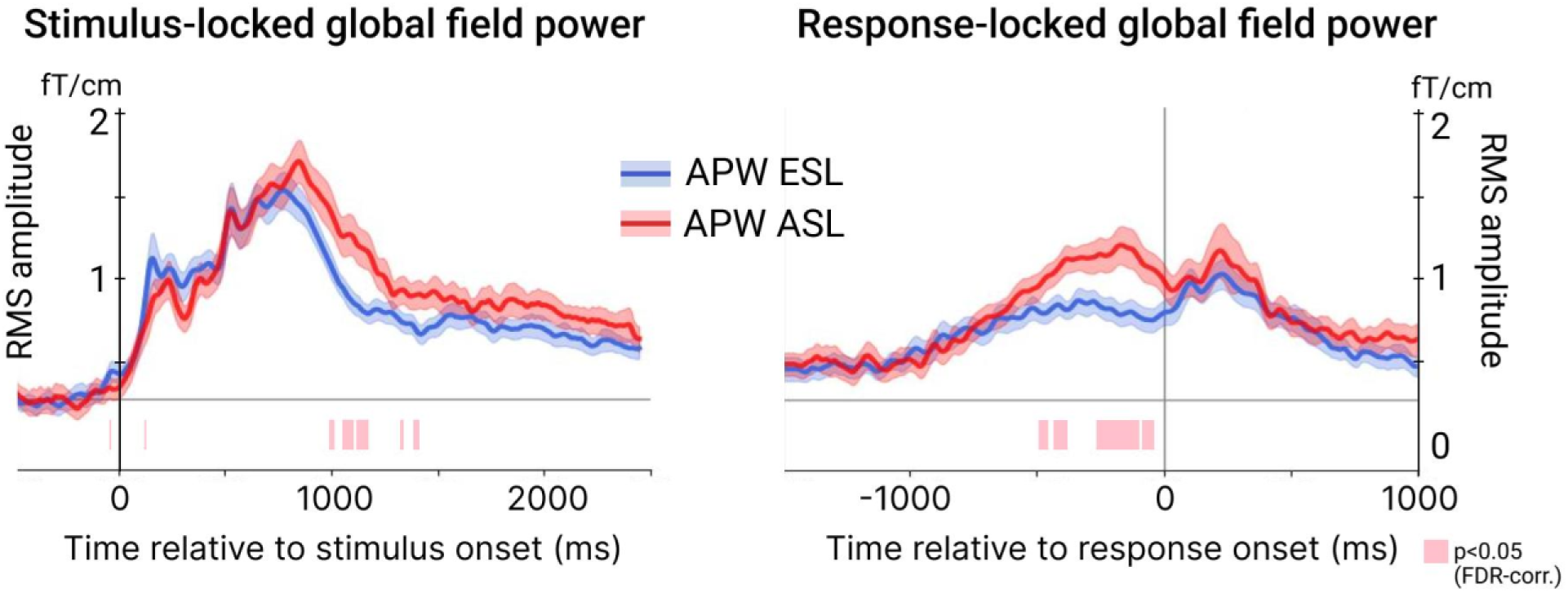

Learning effect in trials matched for the response times. The time courses of the global field power at ESL and ASL on the left are aligned to the stimulus onset, the time courses on the right - to the onset of the motor response. The shaded area around a time course represents the standard error of the mean (SEM). The pink horizontal bars under time courses represent time intervals with significant between-condition differences (p<.05, FDR-corr.).

## References

Aglioti, S. M., & Pazzaglia, M. (2010). Representing actions through their sound. Experimental Brain Research, 206(2), 141–151. 10.1007/s00221-010-2344-x

Aminoff, E. M., Kveraga, K., & Bar, M. (2013). The role of the parahippocampal cortex in cognition. Trends in Cognitive Sciences, 17(8), 379–390. 10.1016/j.tics.2013.06.009

Bangert, M., Peschel, T., Schlaug, G., Rotte, M., Drescher, D., Hinrichs, H.,…Altenmüller, E. (2006). Shared networks for auditory and motor processing in professional pianists: Evidence from fMRI conjunction. NeuroImage, 30(3), 917–926. 10.1016/j.neuroimage.2005.10.044

Barsalou, L. (2003). Situated simulation in the human conceptual system. Language and Cognitive Processes, 18(5–6), 513–562. 10.1080/01690960344000026

Bates, D. (2010). lme4: Linear mixed-effects models using S4 classes. R package version 0.999375–33. Http://CRAN.R-Project.Org/Package=Lme4.

Battaglia, F. P., Benchenane, K., Sirota, A., Pennartz, C. M. A., & Wiener, S. I. (2011). The hippocampus: hub of brain network communication for memory. Trends in Cognitive Sciences, 15(7), 310–318. 10.1016/j.tics.2011.05.008

Benjamini, Y., & Hochberg, Y. (1995). Controlling the False Discovery Rate: A Practical and Powerful Approach to Multiple Testing. Journal of the Royal Statistical Society: Series B (Methodological), 57(1), 289–300. 10.1111/j.2517-6161.1995.tb02031.x

Bi, G., & Poo, M. (1998). Synaptic Modifications in Cultured Hippocampal Neurons: Dependence on Spike Timing, Synaptic Strength, and Postsynaptic Cell Type. The Journal of Neuroscience, 18(24), 10464 LP – 10472. 10.1523/JNEUROSCI.18-24-10464.1998

Bianco, V., Perri, R. L., Berchicci, M., Quinzi, F., Spinelli, D., & Di Russo, F. (2020). Modality-specific sensory readiness for upcoming events revealed by slow cortical potentials. Brain Structure and Function, 225(1), 149–159. 10.1007/s00429-019-01993-8

Brunia, C H M, & van Boxtel, G. J. M. (2004). Anticipatory attention to verbal and non-verbal stimuli is reflected in a modality-specific SPN. Experimental Brain Research, 156(2), 231–239. 10.1007/s00221-003-1780-2

Brunia, Cornelis H M, van Boxtel, G. J. M., & Böcker, K. B. E. (2011). Negative Slow Waves as Indices of Anticipation: The Bereitschaftspotential, the Contingent Negative Variation, and the Stimulus-Preceding Negativity (E. S. Kappenman & S. J. Luck, Eds.). The Oxford Handbook of Event-Related Potential Components, p. 0. 10.1093/oxfordhb/9780195374148.013.0108

Burgess, J. D., Lum, J. A. G., Hohwy, J., & Enticott, P. G. (2017). Echoes on the motor network: how internal motor control structures afford sensory experience. Brain Structure and Function, 222(9), 3865–3888. 10.1007/s00429-017-1484-1

Cheyne, D., Bakhtazad, L., & Gaetz, W. (2006). Spatiotemporal mapping of cortical activity accompanying voluntary movements using an event-related beamforming approach. Human Brain Mapping, 27(3), 213–229. 10.1002/hbm.20178

Dan, Y., & Poo, M.-M. (2006). Spike Timing-Dependent Plasticity: From Synapse to Perception. Physiological Reviews, 86(3), 1033–1048. 10.1152/physrev.00030.2005

Destrieux, C., Fischl, B., Dale, A., & Halgren, E. (2010). Automatic parcellation of human cortical gyri and sulci using standard anatomical nomenclature. NeuroImage, 53(1), 1–15. 10.1016/j.neuroimage.2010.06.010

Drew, P. J., & Abbott, L. F. (2006). Extending the effects of spike-timing-dependent plasticity to behavioral timescales. Proceedings of the National Academy of Sciences, 103(23), 8876–8881. 10.1073/pnas.0600676103

Egger, V., Feldmeyer, D., & Sakmann, B. (1999). Coincidence detection and changes of synaptic efficacy in spiny stellate neurons in rat barrel cortex. Nature Neuroscience, 2(12), 1098–1105. 10.1038/16026

Erlbeck, H., Kübler, A., Kotchoubey, B., & Veser, S. (2014). Task instructions modulate the attentional mode affecting the auditory MMN and the semantic N400. Frontiers in Human Neuroscience, 8. Retrieved from https://www.frontiersin.org/journals/human-neuroscience/articles/10.3389/fnhum.2014.00654

Fan, Z., Muthukumaraswamy, S. D., Singh, K. D., & Shapiro, K. (2012). The role of sustained posterior brain activity in the serial chaining of two cognitive operations: A MEG study. Psychophysiology, 49(8), 1133–1144. 10.1111/j.1469-8986.2012.01391.x

Gallistel, C. R., Fairhurst, S., & Balsam, P. (2004). The learning curve: Implications of a quantitative analysis. Proceedings of the National Academy of Sciences, 101(36), 13124–13131. 10.1073/pnas.0404965101

Garcés, P., López-Sanz, D., Maestú, F., & Pereda, E. (2017). Choice of Magnetometers and Gradiometers after Signal Space Separation. Sensors, Vol. 17. 10.3390/s17122926

Gebel, B., Braun, C., Kaza, E., Altenmüller, E., & Lotze, M. (2013). Instrument specific brain activation in sensorimotor and auditory representation in musicians. NeuroImage, 74, 37–44. 10.1016/j.neuroimage.2013.02.021

Gramfort, A., Luessi, M., Larson, E., Engemann, D. A., Strohmeier, D., Brodbeck, C.,…Hämäläinen, M. S. (2014). MNE software for processing MEG and EEG data. NeuroImage, 86, 446–460. 10.1016/j.neuroimage.2013.10.027

Gross, J., Baillet, S., Barnes, G. R., Henson, R. N., Hillebrand, A., Jensen, O.,…Schoffelen, J.-M. (2013). Good practice for conducting and reporting MEG research. NeuroImage, 65, 349–363. 10.1016/j.neuroimage.2012.10.001

Günseli, E., Fahrenfort, J. J., van Moorselaar, D., Daoultzis, K. C., Meeter, M., & Olivers, C. N. L. (2019). EEG dynamics reveal a dissociation between storage and selective attention within working memory. Scientific Reports, 9(1), 13499. 10.1038/s41598-019-49577-0

Hämäläinen, M., Hari, R., Ilmoniemi, R. J., Knuutila, J., & Lounasmaa, O. V. (1993). Magnetoencephalography---theory, instrumentation, and applications to noninvasive studies of the working human brain. Reviews of Modern Physics, 65(2), 413–497. 10.1103/RevModPhys.65.413

Hebb, D. O. (1949). The first stage of perception: growth of the assembly. The Organization of Behavior, 4(60), 60–78.

Hultén, A., Schoffelen, J.-M., Uddén, J., Lam, N. H. L., & Hagoort, P. (2019). How the brain makes sense beyond the processing of single words – An MEG study. NeuroImage, 186, 586–594. 10.1016/j.neuroimage.2018.11.035

Jafarpour, A., Penny, W., Barnes, G., Knight, R. T., & Duzel, E. (2017). Working Memory Replay Prioritizes Weakly Attended Events. Eneuro, 4(4), ENEURO.0171-17.2017. 10.1523/ENEURO.0171-17.2017

Jefferies, E. (2013). The neural basis of semantic cognition: Converging evidence from neuropsychology, neuroimaging and TMS. Cortex, 49(3), 611–625. 10.1016/j.cortex.2012.10.008

Kaplan, J. T., & Iacoboni, M. (2007). Multimodal action representation in human left ventral premotor cortex. Cognitive Processing, 8(2), 103–113. 10.1007/s10339-007-0165-z

Koch, G., Ponzo, V., Di Lorenzo, F., Caltagirone, C., & Veniero, D. (2013). Hebbian and Anti-Hebbian Spike-Timing-Dependent Plasticity of Human Cortico-Cortical Connections. The Journal of Neuroscience, 33(23), 9725 LP – 9733. 10.1523/JNEUROSCI.4988-12.2013

Kristeva-Feige, R., Rossi, S., Feige, B., Mergner, T., Lücking, C. H., & Rossini, P. M. (1997). The bereitschaftspotential paradigm in investigating voluntary movement organization in humans using magnetoencephalography (MEG) Brain Research Protocols, 1(1), 13–22. 10.1016/S1385-299X(97)80327-3

Kuo, B.-C., Nobre, A. C., Scerif, G., & Astle, D. E. (2016). Top–Down Activation of Spatiotopic Sensory Codes in Perceptual and Working Memory Search. Journal of Cognitive Neuroscience, 28(7), 996–1009. 10.1162/jocn_a_00952

Kutas, M., & Federmeier, K. D. (2011). Thirty years and counting: Finding meaning in the N400 component of the event related brain potential (ERP). Annual Review of Psychology, 62, 621.

León-Cabrera, P., Hjortdal, A., Berthelsen, S. G., Rodríguez-Fornells, A., & Roll, M. (2024). Neurophysiological signatures of prediction in language: A critical review of anticipatory negativities. Neuroscience & Biobehavioral Reviews, 160, 105624. 10.1016/j.neubiorev.2024.105624

Lewis-Peacock, J. A., Drysdale, A. T., Oberauer, K., & Postle, B. R. (2012). Neural Evidence for a Distinction between Short-term Memory and the Focus of Attention. Journal of Cognitive Neuroscience, 24(1), 61–79. 10.1162/jocn_a_00140

Logan, G. D. (1979). On the use of a concurrent memory load to measure attention and automaticity. Journal of Experimental Psychology: Human Perception and Performance, 5(2), 189–207. 10.1037/0096-1523.5.2.189

Maess, B., Herrmann, C. S., Hahne, A., Nakamura, A., & Friederici, A. D. (2006). Localizing the distributed language network responsible for the N400 measured by MEG during auditory sentence processing. Brain Research, 1096(1), 163–172. 10.1016/j.brainres.2006.04.037

Masse, N. Y., Rosen, M. C., & Freedman, D. J. (2020). Reevaluating the Role of Persistent Neural Activity in Short-Term Memory. Trends in Cognitive Sciences, 24(3), 242–258. 10.1016/j.tics.2019.12.014

McCarthy, G., & Nobre, A. C. (1993). Modulation of semantic processing by spatial selective attention. Electroencephalography and Clinical Neurophysiology/Evoked Potentials Section, 88(3), 210–219. 10.1016/0168-5597(93)90005-A

Michelmann, S., Bowman, H., & Hanslmayr, S. (2018). Replay of Stimulus-specific Temporal Patterns during Associative Memory Formation. Journal of Cognitive Neuroscience, 30(11), 1577–1589. 10.1162/jocn_a_01304

Neuringer, A. (2002). Operant variability: Evidence, functions, and theory. Psychonomic Bulletin & Review, 9(4), 672–705. 10.3758/BF03196324

Nicoll, R. A. (2017). A Brief History of Long-Term Potentiation. Neuron, 93(2), 281–290. 10.1016/j.neuron.2016.12.015

O’Rourke, T. B., & Holcomb, P. J. (2002). Electrophysiological evidence for the efficiency of spoken word processing. Biological Psychology, 60(2), 121–150. 10.1016/S0301-0511(02)00045-5

Ohgami, Y., Kotani, Y., Hiraku, S., Aihara, Y., & Ishii, M. (2004). Effects of reward and stimulus modality on stimulus-preceding negativity. Psychophysiology, 41(5), 729–738. 10.1111/j.1469-8986.2004.00203.x

Oldfield, R. C. (1971). The assessment and analysis of handedness: The Edinburgh inventory. Neuropsychologia, 9(1), 97–113. 10.1016/0028-3932(71)90067-4

Poeppel, D., Idsardi, W. J., & van Wassenhove, V. (2007). Speech perception at the interface of neurobiology and linguistics. Philosophical Transactions of the Royal Society B: Biological Sciences, 363(1493), 1071–1086. 10.1098/rstb.2007.2160

Praamstra, P., Schmitz, F., Freund, H.-J., & Schnitzler, A. (1999). Magneto-encephalographic correlates of the lateralized readiness potential. Cognitive Brain Research, 8(2), 77–85. 10.1016/S0926-6410(99)00008-7

Pulvermüller, F. (2005). Brain mechanisms linking language and action. Nature Reviews Neuroscience, 6, 576. Retrieved from 10.1038/nrn1706

Pulvermüller, F. (2018). Neural reuse of action perception circuits for language, concepts and communication. Progress in Neurobiology, 160, 1–44. 10.1016/j.pneurobio.2017.07.001

Razorenova, A. M., Chernyshev, B. V, Nikolaeva, A. Y., Butorina, A. V, Prokofyev, A. O., Tyulenev, N. B., & Stroganova, T. A. (2020). Rapid Cortical Plasticity Induced by Active Associative Learning of Novel Words in Human Adults. Frontiers in Neuroscience, Vol. 14, p. 895. Retrieved from https://www.frontiersin.org/article/10.3389/fnins.2020.00895

Rolls, E. T., Deco, G., Huang, C.-C., & Feng, J. (2022). The effective connectivity of the human hippocampal memory system. Cerebral Cortex, 32(17), 3706–3725. 10.1093/cercor/bhab442

Schurger, A., Hu, P. “Ben,” Pak, J., & Roskies, A. L. (2021). What Is the Readiness Potential? Trends in Cognitive Sciences, 25(7), 558–570. 10.1016/j.tics.2021.04.001

Shtyrov, Y., Butorina, A., Nikolaeva, A., & Stroganova, T. (2014). Automatic ultrarapid activation and inhibition of cortical motor systems in spoken word comprehension. Proceedings of the National Academy of Sciences of the United States of America, 111(18), E1918--23. 10.1073/pnas.1323158111

Shu, Y., Hasenstaub, A., & McCormick, D. A. (2003). Turning on and off recurrent balanced cortical activity. Nature, 423(6937), 288–293. 10.1038/nature01616

Silva Pereira, S., Hindriks, R., Mühlberg, S., Maris, E., van Ede, F., Griffa, A.,…Deco, G. (2017). Effect of Field Spread on Resting-State Magneto Encephalography Functional Network Analysis: A Computational Modeling Study. Brain Connectivity, 7(9), 541–557. 10.1089/brain.2017.0525

Sutherland, N. S., & Mackintosh, N. J. (1971). Mechanisms of animal discrimination learning. Academic Press.

Tandonnet, C., Burle, B., Vidal, F., & Hasbroucq, T. (2003). The influence of time preparation on motor processes assessed by surface Laplacian estimation. Clinical Neurophysiology, 114(12), 2376–2384. 10.1016/S1388-2457(03)00253-0

Taulu, S., Simola, J., & Kajola, M. (2005). Applications of the Signal Space Separation Method. IEEE Transactions on Signal Processing, Vol. 53, pp. 3359–3372. 10.1109/TSP.2005.853302

Teyler, T. J., & DiScenna, P. (1985). The role of hippocampus in memory: A hypothesis. Neuroscience & Biobehavioral Reviews, 9(3), 377–389. 10.1016/0149-7634(85)90016-8

Teyler, T. J., & Rudy, J. W. (2007). The hippocampal indexing theory and episodic memory: Updating the index. Hippocampus, 17(12), 1158–1169. 10.1002/hipo.20350

Tomasello, R., Garagnani, M., Wennekers, T., & Pulvermüller, F. (2017). Brain connections of words, perceptions and actions: A neurobiological model of spatio-temporal semantic activation in the human cortex. Neuropsychologia, 98, 111–129. 10.1016/j.neuropsychologia.2016.07.004

Turkeltaub, P. E., & Branch Coslett, H. (2010). Localization of sublexical speech perception components. Brain and Language, 114(1), 1–15. 10.1016/j.bandl.2010.03.008

Vogel, E. K., & Machizawa, M. G. (2004). Neural activity predicts individual differences in visual working memory capacity. Nature, 428(6984), 748–751. 10.1038/nature02447

Vrba, J., & Robinson, S. E. (2001). Signal Processing in Magnetoencephalography. Methods, 25(2), 249–271. 10.1006/meth.2001.1238

Wang, J. X., Rogers, L. M., Gross, E. Z., Ryals, A. J., Dokucu, M. E., Brandstatt, K. L.,…Voss, J. L. (2014). Targeted enhancement of cortical-hippocampal brain networks and associative memory. Science, 345(6200), 1054–1057. 10.1126/science.1252900

Wang, X.-J. (2001). Synaptic reverberation underlying mnemonic persistent activity. Trends in Neurosciences, 24(8), 455–463. 10.1016/S0166-2236(00)01868-3

Yousry, T. A., Schmid, U. D., Alkadhi, H., Schmidt, D., Peraud, A., Buettner, A., & Winkler, P. (1997). Localization of the motor hand area to a knob on the precentral gyrus. A new landmark. Brain, 120(1), 141–157. 10.1093/brain/120.1.141

Zhang, L., Xi, J., Xu, G., Shu, H., Wang, X., & Li, P. (2011). Cortical Dynamics of Acoustic and Phonological Processing in Speech Perception. PLOS ONE, 6(6), e20963. Retrieved from 10.1371/journal.pone.0020963

